# Monocot-specific *miR528* act as the post-transcriptional regulator of strigolactone signaling via *Dwarf 3* in rice

**DOI:** 10.1101/2023.11.06.565764

**Authors:** Sonia Balyan, Deepika Sharma, Shivani Kansal, Vaishali Panwar, Ringyao Jajo, Tonu Angaila Chithung, Saurabh Raghuvanshi

## Abstract

miRNAs are a critical component of regulatory mechanisms involved in plant growth, development, and stress response including phytohormone action. The role of miRNAs in regulating the strigolactone (SL) signaling in plants is still unexplored. Here we report that miR528 impacts the SL signaling via the post-transcriptional regulation of *Dwarf 3* (*OsD3*). Comparative miRNome analysis under drought conditions in flag leaf, inflorescence, and roots uncovered the tissue-biased unique stress response of miR528 in drought-tolerant N22 (down in inflorescence and roots; up in flag leaf). Consequently, its target, *OsD3* which encodes for the F-BOX/LRR-REPEAT MAX2 HOMOLOG and is a critical component of the rice SL signaling pathway exhibit opposite expression to miR528 in all three tissues. Interestingly, *OsD3* is a rice-specific target of miR528. miR528 overexpression plants exhibited early heading, increment in the number of tillers, flag leaf length, and width, number of panicle branches, grain per panicle, effective grains per panicle, seed length, and photosynthesis rate. miR528 overexpression plants exhibited similar traits to *Osd3* mutants including high tiller number, root architecture under low nitrogen conditions, and moderate insensitivity to GR24 and TIS1 levels in addition to shared transcriptional regulation. Furthermore, SL (GR24) negatively impacts the miR528 transcription. The impact of the miR528:D3 module is also reflected in the D53-mediated downstream signaling involving IPA1 regulon.

## Introduction

MicroRNAs (miRNAs) are tiny, 21 to 22 nucleotides (nt) long, ubiquitous crucial transcriptional, post-transcriptional, and translational regulators of eukaryotic gene regulatory networks involved in shaping nearly every aspect of growth and development including response to external stimuli. miRNAs score their presence in every domain of life including plants and animals and shared a congruent framework of their biogenesis, mode of action, and turnover in different lineages with some divergence. The miRNome expansion in plants through exploiting the high-throughput sequencing technologies showed that besides the deeply conserved miRNA families, most of the miRNA families are species or lineage-specific (Cuperus et al., 2011). One such miRNA family is miR528, which is restricted only to class *Liliopsida* and recorded its presence in *Araceae*, *Arecaceae*, *Asparagaceae*, *Musaceae*, *Orchidaceae*, and *Poaceae* (Guo et al. 2019). Several investigations uncovered the multifarious roles of miR528 in monocots. In rice, *miR528* was upregulated in response to aluminum excess (Lima et al., 2011), cadmium (Ding et al., 2011), zinc (Zeng et al. 2019), and arsenite (Liu et al., 2012, Sharma et al., 2014) and downregulated in response to H_2_O_2_ (Li et al., 2011), salt (Barrera-Figueroa et al., 2012) and nitrogen starvation (Nischal et al., 2012). Its levels decrease upon drought in sugarcane (Ferreira et al., 2012) and wheat (Kantar et al., 2011), and nitrogen starvation in maize (Zhao et al., 2012). Interestingly miR528 was induced in water-logging tolerant maize lines and decreased in sensitive lines under short-term water-logging stress (Liu et al., 2012). In wheat, *miR528* was downregulated in heat stress (Ragupathy et al., 2016). *miR528* preferentially target genes encoding copper-containing proteins (Balyan et al., 2017; Zhu et al., 2020) and function as the critical modulator of cellular reactive oxygen species (ROS) homeostasis. Recently, a study characterized the negative role of *miR528* in the viral resistance of rice by cleaving L-ascorbate oxidase mRNA (Wu et al., 2017). In addition, the study showed that AGO18 sequesters the *miR528* in response to viral infection thereby decreasing the active *miR528* for the cleavage of *ascorbate oxidase* (*AO*), leading to higher basal reactive oxygen species (Wu et al., 2017). Recently, miR528 shown to be implicated in rice pollen intine formation by directly regulating uclacyanin family member (*OsUCL23)* (Zhang et al., 2020). In banana, under cold stress, *miR528* was downregulated and its target polyphenol oxidase (PPO) was upregulated leading to ROS surge and peel browning of its fruit (Zhu et al., 2020). Besides the canonical copper-binding targets, miR528 regulates the other species-specific target genes involved in diverse metabolic pathways. One such validated target of miR528 is *OsD3* (F-box protein), a critical component in strigolactone signaling in rice. Strigolactones are recently emerged as the class of plant hormones of utmost agricultural significance due to their involvement in the regulation of plant-rhizosphere parasitic as well as symbiotic interactions and several traits determining crop yields and biomass including root and shoot architecture and nutrient and defense mechanisms (Akiyama et al., 2005; Bürger & Chory, 2020a; Domagalska & Leyser, 2011; Gomez-Roldan et al., 2008; Yan et al., 1998). Although *OsD3* is one of the high-confidence targets of *miR528* but still the implications of the *miR528:D3* module in shaping rice SL signaling were unclear. The present study uncovers the novel role of miR528 in regulating the SL signaling and its downstream transcriptional regulations involving the *dwarf 53* and Ideal Plant Architecture 1 (D53:IPA1) regulon-mediated processes. Further, the study delineated that SL derives the negative transcriptional regulation of miR528.

## Results

### Drought-regulated miRNome of the root of N22

The function of roots is indispensable in determining the tolerance of plants to drought. Roots are the first organs to sense and orchestrate the defense response against water stress to ensure plant survival (Ahmadi et al., 2014). Nagina 22 is one of the most important drought and heat-tolerant rice cultivar (Selote et al. 2004; Prasad et al. 2006; Jagadish et al. 2007) and served as a tolerant donor in several breeding studies. Several studies untapped its genome-wide transcriptome in different tissues and stress regimes (Gour et al., 2021; Negi et al., 2016; Sevanthi et al., 2021) but the root-specific miRNome is still overlooked. Therefore, the present study delineated the role of drought-regulated miRNome in roots of N22 under field conditions. As a result, a total of 272 and 259 miRNAs were identified in control and drought-stressed roots respectively (Table S1). Drought leads to the upregulation of 24 miRNAs while 14 exhibited downregulation in response to drought (Figure 1a). *miR1320-5p, miR1432-5p, miR156a-j/k/l-5p, miR164e, miR1846e, miR1850.1, miR1861b/f/l, miR319a-3p, miR3979-3p, miR5144-5p, miR528-5p and miR535-5p* were downregulated in roots (Table S2). While four miR156 family member (*b-3p, c-3p/g-3p, f-3p/h-3p/l-3p, j-3p*), *miR159a.1/b, miR160a-d-5p, miR167h-3p, miR168a-3p, miR171h, miR1862d, miR1862e, miR1871, miR1876, miR2878-5p, miR393b-3p, miR396a/b-5p, miR437, miR530-5p, miR5802, miR812* members (a-e, p, s, v) and miR818a-e were upregulated in roots (Table S2). The target genes of differentially regulated miRNAs were obtained from the degradome data analysis published in our previous study (Balyan et al., 2017). The degradome-identified targets of drought-regulated miRNAs were analyzed for enrichment of gene ontology and trait ontology terms using ShinyGO (http://bioinformatics.sdstate.edu/go/) and oryzabase (https://shigen.nig.ac.jp/rice/oryzabase/locale/change?lang=en). Several targets were observed to be involved in growth, yield as well stress-associated traits, and GO terms (Figure S1). Further 23 miRNA-target modules are part of rice metabolic pathways (Table S3). To get more insights into the role of miRNAs in rice drought regulation, the root miRNome was compared with the miRNome of flag leaf and spikelets in N22 (Balyan et al., 2017; Figure 1b). As a result, 7 miRNAs (*miR3979-3p, miR156b-3p, miR3979-5p, miR156f/h/l-3p, miR1320-3p, miR156c-3p/g-3p, and miR166a-5p/e-5p*) exhibited enriched expression (≥5 fold) in roots as compared to flag leaf and spikelet (Figure 1c). *miR3979-3p* and *miR156f/h/l-3p* represent the root-specific drought-regulated miRNA. Further, when the expression profile of 38 drought-regulated miRNAs identified in roots was compared with flag leaf and spikelets (Balyan et al., 2017), only four miRNAs (*miR528-5p, miR812a-e, miR812v, and miR5802*) were regulated in all three tissues under drought in N22 (Figure 1d-e). *miR528-5p* followed drought-mediated downregulation in spikelets and roots while upregulated in flag leaf. While *miR812a-e*, *miR812v*, and *miR5802* exhibited upregulation in roots and downregulated in flag leaf and spikelets. Furthermore, miR528-5p exhibited inverse expression in the flag leaf of drought-sensitive and tolerant cultivars suggesting its critical function in tissue-mediated drought regulation (Figure 1d). Therefore, *miR528* was selected for detailed characterization in rice.

**Figure 1.**
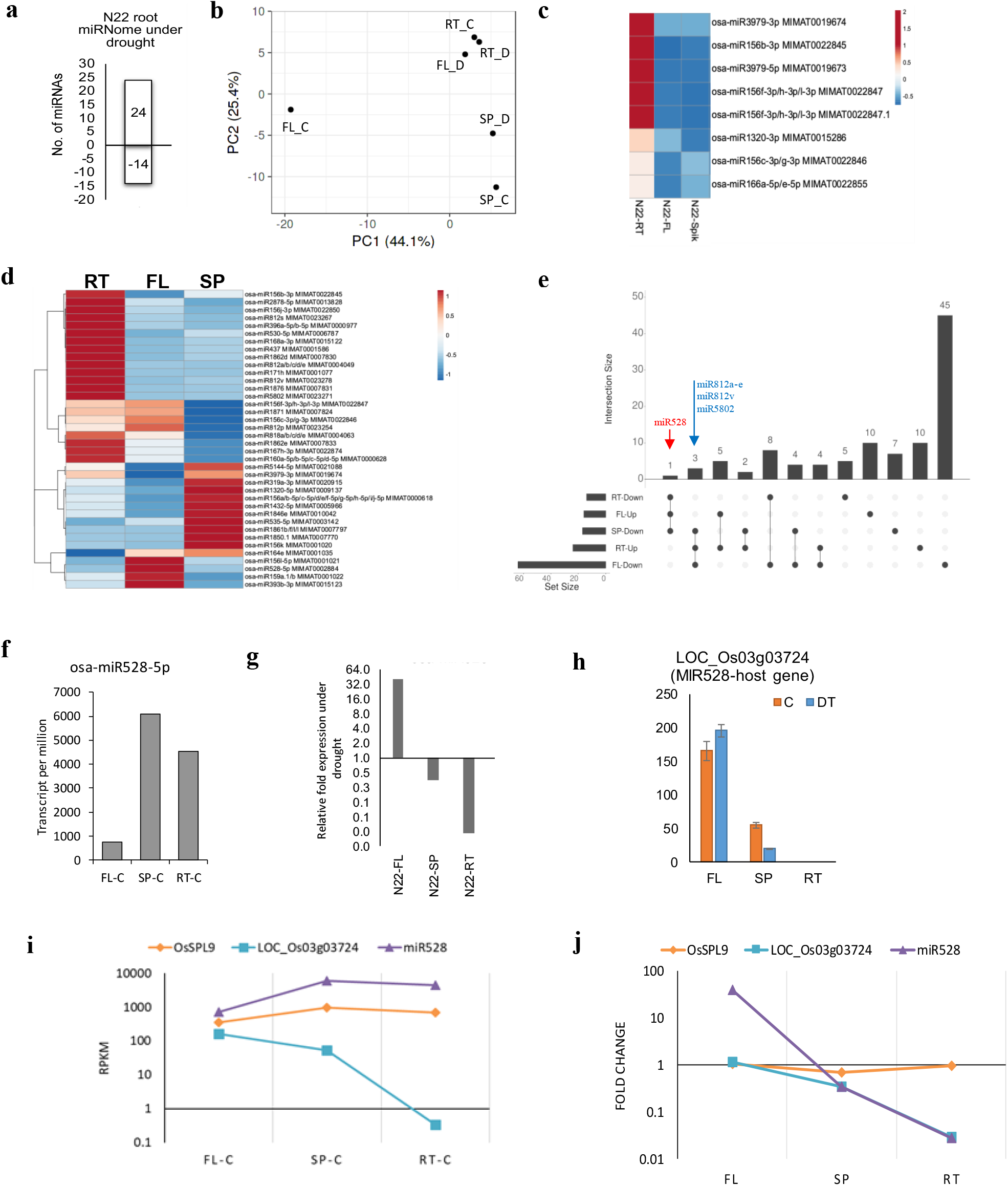
miR528 exhibit tissue-mediated drought response in N22. **(a)** Number of upregulated (FC≥2) and downregulated (FC≤-2) miRNAs identified by comparative miRNome analysis of roots under control and drought stress conditions. **(b)** The Principal Component Analysis plot depicting the comparison of different tissue-specific (RT=root; FL=flag leaf; SP= spikelets) miRNome of N22 under control and drought conditions. **(c)** Heat map of root-enriched miRNAs in rice cultivars (N22 and PB1). The miRNA expression levels of root were compared with flag leaf and spikelet of N22 and PB1 (Balyan et al. 2017) and miRNA with ≥ 5 fold higher expression in root as compared to cumulative expression in flag leaf and spikelet were shown in the heat map. **(d)** Drought regulated miRNAs identified in roots and their comparative expression in flag leaf and inflorescence. **(e)** The upset plots show the interaction of differentially regulated miRNAs in different tissue of N22 in response to drought stress. The miRNAs exhibiting the drought regulation in all three tissues were highlighted **(f)** The expression of miR528-5p in FL, SP and RT of N22 **(g)** Drought-regulated expression profile of miR528-5p in FL, SP and RT of N22. **(h)** the expression of host gene (LOC_Os03g03724) of miR528 in FL, SP and RT of N22. **(i-j)** The comparative tissue biased **(i)** and drought regulated **(j)** expression of OsSPL9, miR528-host gene and miR528-5p in N22.

### Monocot-specific miR528 exhibit tissue-biased drought response in N22

*miR528* is a Liliopsida-specific miRNA and annotated in *Araceae, Arecaceae, Asparagaceae, Musaceae, Orchidaceae, Poaceae, and Bromeliaceae* (Figure S2a). To understand the sequence variation among the different species of Liliopsida, the mature and precursor sequences were extracted from different public repositories (miRBase, pmiRen, and sRNanno). The mature sequence of *miR528* (n=41) is highly conserved in the above-mentioned groups and out of 41 precursors, 39 gave rise to major canonical *miR528* (UGGAAGGGGCAUGCAGAGGAG, 21nt) form, while two minor sequence variants i.e. *pda-miR528a*(UGGAAGGG**A**CAUGCAGAGGAA) and *Egu-miR528* (UGGAAGGGGCAUGCAGAGG**C**G) were also reported from two precursors. All miR528 forms are 21nt in length (Figure S2b-c). The sequence alignment of miR528 precursors demonstrates clear conservation peaks in and around miR and miR* region (Figure S2b). Besides the highly conserved mature sequence, the critical bases (GC at position 8-9 and 18-19, marked by *) in *miR528*\* reported for *miR528* precursor processing (Narjala et al., 2020) are well conserved in plants (Figure S2b). Also, the position of two mismatches at the 12^th^ and 16^th^ positions are consistent in all monocots but with variation in base (Figure S2b). The phylogenetic analysis using maximum likelihood divided the *miR528* family into three major clades C-I to III. Oryza clustered together with *Phyllostachys* in C-I. While other Poales forms a major cluster C-II together. Other monocots belonging to Araceae, Arecaceae, Asparagaceae, Musaceae, Orchidaceae, and Bromeliaceae are grouped in C-III.

In rice, *miR528* is exonic miRNA and its locus is located in 5′ UTR of LOC_Os03g03724 (expressed protein). To understand the function of *miR528*, its tissue, and drought-regulated expression were comprehensively examined in N22. Under control conditions, *miR528* was highly abundant in spikelet and root as compared to flag leaf (Figure 1f). But under drought conditions, *miR528* exhibits upregulation in flag leaf while downregulation in spikelet and roots (Figure 1g). This suggests that the *miR528* is governed by tissue as well as drought-mediated regulations in N22. In alignment with the miR528, its host gene also showed increased levels in flag leaf but downregulation in spikelet, while the host gene remains undetected in roots (Figure 1h). In summary *miR528* shares parallel drought regulation with its host gene. Interestingly the tissue-mediated expression dynamics between *miR528* and its host are opposite to each other, suggesting the existence of an additional regulatory layer at the level of its transcription in rice (Figure 1i). *miR528* is regulated by SPL9 in rice, which is the master regulator of copper homeostasis (Balyan et al., 2017; Yao et al., 2019). Tissue-biased regulation of *miR528* was similar to *SPL9* while the opposite pattern was observed for the host gene. While the *SPL9* is maintained in flag leaf and roots but exhibited slight downregulation in spikelet while *miR528* and its host gene followed similar drought regulation (Figure 1i-j). To further enquire into the role of epigenetic regulation in deriving the transcription of *miR528*, the whole genome bisulfite sequencing of N22 under control as well as drought conditions in corresponding tissues was used to extract the methylation signature around its host gene. Neither the *miR528* host nor its promoter showed methylation signatures suggesting that the *miR528* locus is not epigenetically regulated in N22 (Figure S3). The upstream methylation signatures lie within the gene body of another upstream gene beyond the 1 kb promoter region.

### *miR528* is the critical post-transcriptional regulator of rice transcriptome

As miRNAs exert their function through the post-transcriptional regulation of their target mRNAs, therefore it is critical to study the target pool of *miR528* to delineate the roles played by *miR528* in rice. The targets of *miR528* were predicted by psRNATarget (score ≤ 5) using cDNA of rice (http://rice.uga.edu/pub/data/Eukaryotic_Projects/o_sativa/annotation_dbs/pseudomolecules/version_7.0/all.dir/) and mature *miR528* as inputs. Further, the targets having degradome support were extracted from pmiRen (https://www.pmiren.com/), tarbase (https://dianalab.e-ce.uth.gr/html/diana/web/index.php?r=tarbasev8), and a study by Balyan et al. 2017. A total of 132 target genes were predicted as miR528 targets and 19 genes were validated with degradome support. 4 canonical high-confidence target genes including LOC_Os06g06050 (*OsD3*), LOC_Os06g37150 (*OsAAO2*), LOC_Os08g04310 (*OsUCL23*), and LOC_Os07g38290 (plastocyanin-like domain-containing protein) were identified by all sources (Table S4). But still, some targets may be uniquely regulated in a specific tissue or environmental condition and therefore skipped by degradome data.

To predict the possible regulatory roles of miR528 via post-transcriptional regulation of its targets, the GO enrichment analysis of target genes was performed using ShinyGO (http://bioinformatics.sdstate.edu/go/) (Figure S4a). The results highlighted the enrichment of processes involving the copper-requiring proteins including ascorbate oxidase, multicopper oxidases, and phytocyanins in addition to terms associated with brassinosteroids biosynthetic processes. Furthermore, the trait ontology analysis of targets overshines a few critical regulators of rice growth and stress response (Figure S4b). 12 genes encoding for copper-requiring proteins form a sub-network associated with several terms associated with rice metabolic pathways. Importantly, two targets {LOC_Os06g06050 (OsD3) and LOC_Os07g01240 (SRL1)} seem to be critical factors for plant growth and stress response as evidenced by their association with > 20 GO and TO terms (Figure S4b). Furthermore, several targets need to be functionally characterized to completely delineate the function and implication of miR528/target node. Considering the limited degradome data regarding the tissue, stress conditions, and cultivar backgrounds, it’s possible that several genuine targets may be overlooked. Therefore, to analyze the complete miR528-target module in N22, the miR528 expression was compared with the target genes in terms of tissue enrichment and drought stress response. Comparison of expression fold change (RT/FL) from root to flag leaf demonstrated that 45 targets followed higher expression in root parallel to miR528 while 32 exhibited decreased expression in roots (Figure 2a). Further, when the miR528-target module is compared between inflorescence and flag leaf, 59 targets shared similar expression to miR528 while only 16 exhibited the opposite trend (Figure 2b). When the drought-regulated dynamics of the miR528-target module were analyzed in different tissues, target genes followed unique to shared inverse regulon to miR528 in different tissues of rice. For instance, only two target genes {LOC_Os02g55600 expressed protein; score = 5; predicted only) and LOC_Os06g06050 (*OsD3;* score = 2; high confidence degradome identified)} followed inverse expression pattern to miR528 in all three tissues under drought (Figure 2c-f). While expression of 20 genes showed anti-correlation to miR528 in any two tissues (Figure 2c-f). Moreover, the opposite regulation was uniquely observed for 6, 9, and 39 targets in spikelet, flag leaf, and roots respectively under drought stress conditions (Figure 2f). The above results suggest that the transcriptional regulation of miR528/target pool was dynamically influenced by the tissue type and stress regimen and *OsD3* is the most genuine target in N22.

**Figure 2.**
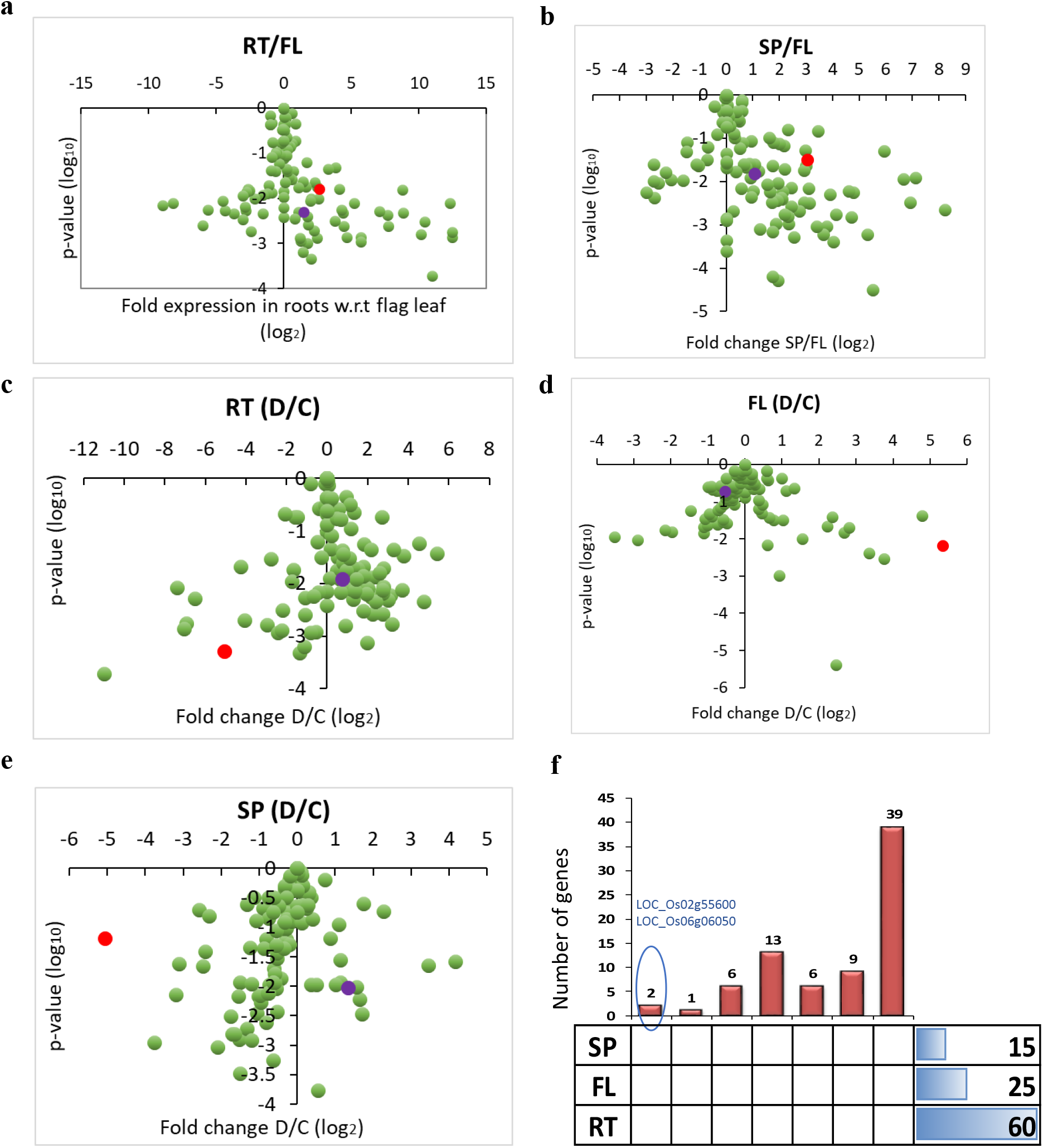
Comparative expression analysis of miR528 and its targets genes in different tissue under drought stress in N22. **(a-e)** The fold expression of miR528-5p (red dot) and its target genes (green dots) is plotted against p-value (log10) between different tissue contexts **(RT/FL:** Root vs Flag Leaf; **SP/FL:** Spikelet vs Flag leaf, **a-b)** and drought response in root (**RT**; **c**), flag leaf (**FL**; **d**) and spikelet (**SP**; **e**). **(f)** Upset plots showing the intersection of targets following anti-correlation in expression to that of miR528-5p in different tissues under drought. SP=spikelet; FL=flag leaf and RT=root. D/C in **c-e** signifies drought vs control.

### Comparative analysis of miR528 sequence variations and haplotype evaluation in rice

Further to identify the variation in miR528 precursor and its upstream cis-regulatory region among the 4726 rice accessions, RiceVarMap2 (http://ricevarmap.ncpgr.cn; Zhao et al. 2021) database was used. The miR528 host gene (LOC_Os03g03724) along with 1Kb upstream cis-regulatory region (Ch03:1,666,247-1,668,963) was screened for sequence variation (SNP and Indels) across 4726 rice accessions. As a result, a total of 32 sequence variants were detected, including 26 in the promoter region and only 6 lies in the gene (Figure S5a). While the region containing the miR528 precursor was almost conserved in rice accession except at position 1667386 (Figure S5a). The variants that lie in the promoter region were used to evaluate the haplotype using the Haplotype Network Analysis tool of RiceVarMap2. As a result, six haplotypes were calculated including haplotypes hap-II, III, IV, V, VI which predominate in indica-accessions (Figure S5b-c). The variants detected in the promoter region may modify the cis-regulatory motifs of any transcription factor. Therefore, the intersection and overlap of the above-detected variants were mapped onto the cis-regulatory motifs identified in the 1Kb promoter of miR528 using New Place database (https://www.dna.affrc.go.jp/PLACE/?action=newplace, Higo et al. 1999). Only six variants coincide with the cis-regulatory sites eg TATABOX, POLASIGI, NTBBFARROLB, S1FBOXSORPS1L21, CAATBOX, and Preconscrhsp70A (Figure S5d).

### miR528 overexpression regulates multiple traits in plants

To excavate the function of miR528 in rice, 88bp precursor of miR528 was PCR amplified from the genomic DNA isolated from the flag leaf of N22 (Figure S6a). The desired fragment was purified and digested with KpnI and SacI and cloned in pB4NU binary vector (Figure S6b-e). Here the expression of MIR528 is driven by the maize ubiquitin promoter. The construct was used to raise the rice transgenic plants over-expressing MIR528 (Figures S6 and 3). The transgenic nature was confirmed by PCR amplification of ubiquitin (Figure S6e). Further, the expression level of miR528-5p was analyzed in different lines at the 2-month-old stage wherein miR528-5p was found to over-express by 3 to 30-fold higher than the wild-type plants (Figure S6f). The target genes were also downregulated in MIR528-OE plants confirming the miR528 activity in transgenics (Figure S6g).

In-depth phenotyping analysis of wild-type (WT) and MIR528-OE plants identified several morphological and yield-associated traits that are modulated upon miR528 overexpression (Figure 3 and S7). At the 7-day old seedling stage, the miR528-OE seedlings exhibited more adventitious and seminal roots with slightly decreased shoot length and root length (Figure S7). Although there is no difference in seedling weight (Figure S7). The total chlorophyll content was more in miR528-OE seedlings due to increased chl-a levels (Figure S7f). At the reproductive stage, MIR528-OE plants showed early heading phenotype around 100 DAS compared to 120 DAS observed for WT plants (Figure 3a-c). In addition, the MIR528-OE had a significant (double) increase in the number of tillers, flag leaf length, and width (Figure 3d-h). While the plant height did not seem to be regulated by MIR528 which is also evident from the length comparison of internodes where the change is insignificant from the 2^nd^ to the 5^th^ internode (Figure 3i-k). Moreover, the miR528 seems to be a positive regulator of rice grain yield as shown by the enhanced number of primary branches of panicle, the number of grain/panicle, and a number of effective grains/panicle (Figure 3l-o). Further, the seed length was also slightly increased in overexpression lines (Figure 3r-s). While the grain width was not significantly affected in overexpression plants (Figure 3p-q). The net photosynthesis rate was also increased upon miR528 overexpression while water use efficiency showed a decreased trend (Figure 3t-u). There was a slight increase in the stomatal conductance in the miR528-OE plants (Figure 3v). All these results suggest that miR528 is a positive regulator of early heading, number of tillers, and grain yield in rice.

**Figure 3.**
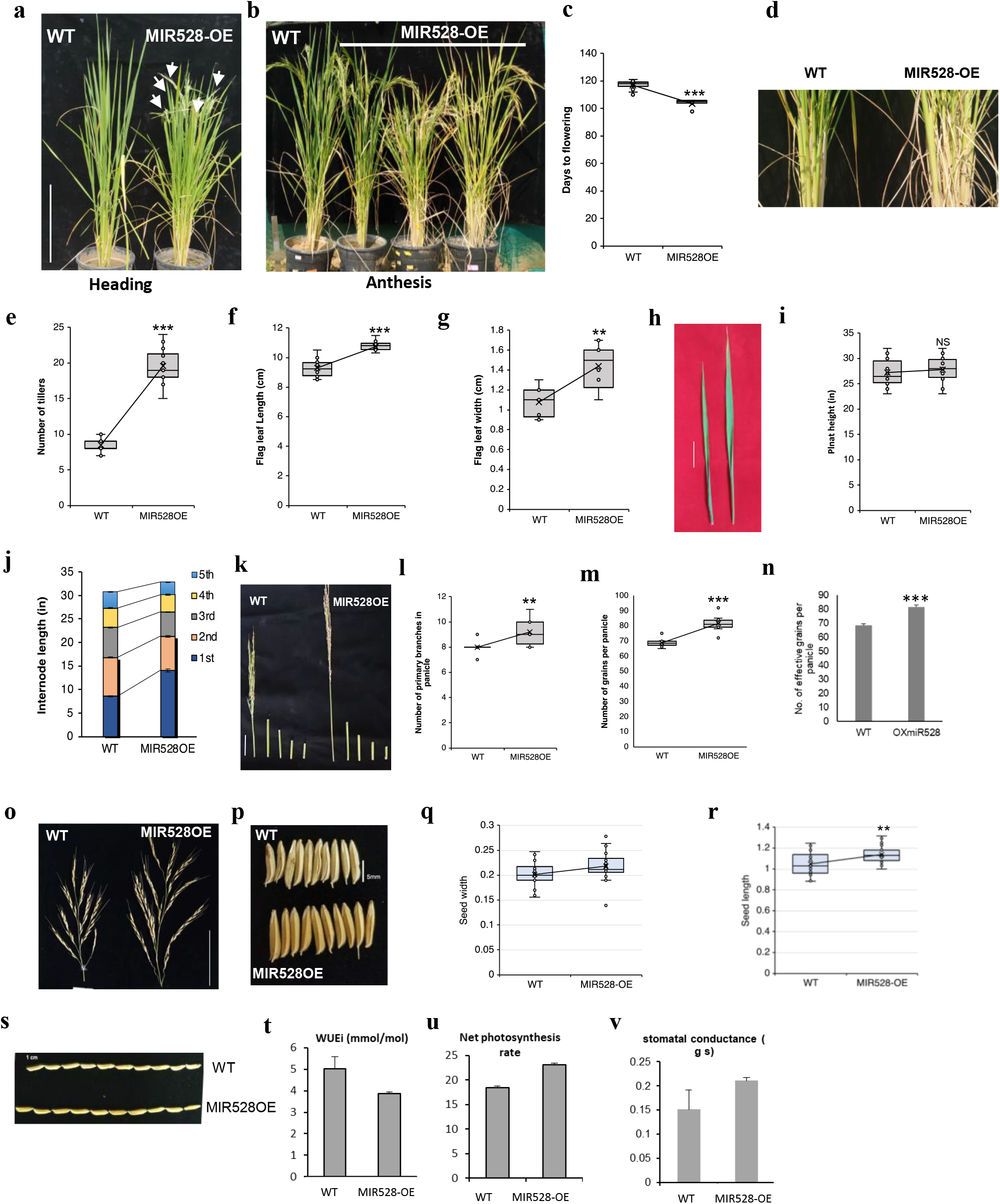
miR528 regulate several important traits in rice. **(a-b)** Phenotypic analysis of wild-type (WT) and MIR528-OE plants at heading **(a)** and anthesis stage **(b).** Arrows points the heading panicles in the MIR528-OE. **(c)** Days to flowering of WT (n=10) and MIR528-OE (n=10) are represented as box plots. **(d)** phenotype showing the number of tillers of WT and miR528-OE plants at heading stage. **(e)** The box plots showing the number of tillers in WT (n=10) and miR528-OE (n=10). **(f-h)** comparison of flag leaf length **(f)**, width **(g)** and phenotype of WT and mI528-OE plants. **(i-v)** Estimation of plant height **(i),** internode length **(j-k),** number of primary branches of panicle **(l)**, number of grains per panicle **(m)**, number of effective grains per panicle **(n-o),** seed length and width **(p-s),** WUS **(t**), net photosynthetic rate **(u)** and stomatal conductance **(v)** of WT and miR528-OE plants. The number of plants used in each quantification are indicated as n and the data points are shown as circles. ***P< 0.001; **P<0.01 and *P<0.05 as per student’s t-test.

### Overexpression of miR528 leads to the transcriptional activation of genes involved in photosynthetic machinery

Next, to enquire into the impact of *miR528* overexpression on the rice transcriptome, the RNA-seq data (paired-end) was generated and analyzed for the shoot tissue of two-week-old WT and MIR528-OE seedlings. The PCA analysis separates the RNA-Seq samples of MIR528-OE and WT (Figure 4a). At first, the transcriptional modulation of miR528 targets was studied to validate the miR528-mediated post-transcriptional regulation of mRNAs. As expected, the target pool of *miR528* was downscaled as shown in the plot (Figure 4b). In addition, RNA-Seq identified 321 genes that are differentially regulated in MIR528-OE shoots including 149 upregulated and 172 downregulated genes (Figure 4c). To further decode the miR528-mediated transcriptional shift, the GO enrichment analysis was performed using ShinyGO (Figure 4d-e). As increased photosynthetic rate was increased in miR528-OE plants, the RNA-seq further confirms the enrichment of genes involved in photosynthesis-associated GO terms *viz* carbon dioxide fixation, photosynthesis light reactions, photosystem II, thylakoid, chloroplast, etc. While the downregulated genes belong to the metallothioneins, oxylipin biosynthesis, metal-ion binding, etc (Figure 4e). Mapping of differentially expressed genes on rice pathways using mapman clearly showed the significant upregulation of several genes involved in light reactions (Figure 4f). Furthermore, the categorization of DEGs according to their associated traits suggests the involvement of miR528 in regulating the salt, drought, light sensitivity, photosynthetic rate and chlorophyll, grain yield, and plant height (Figure 4g).

**Figure 4.**
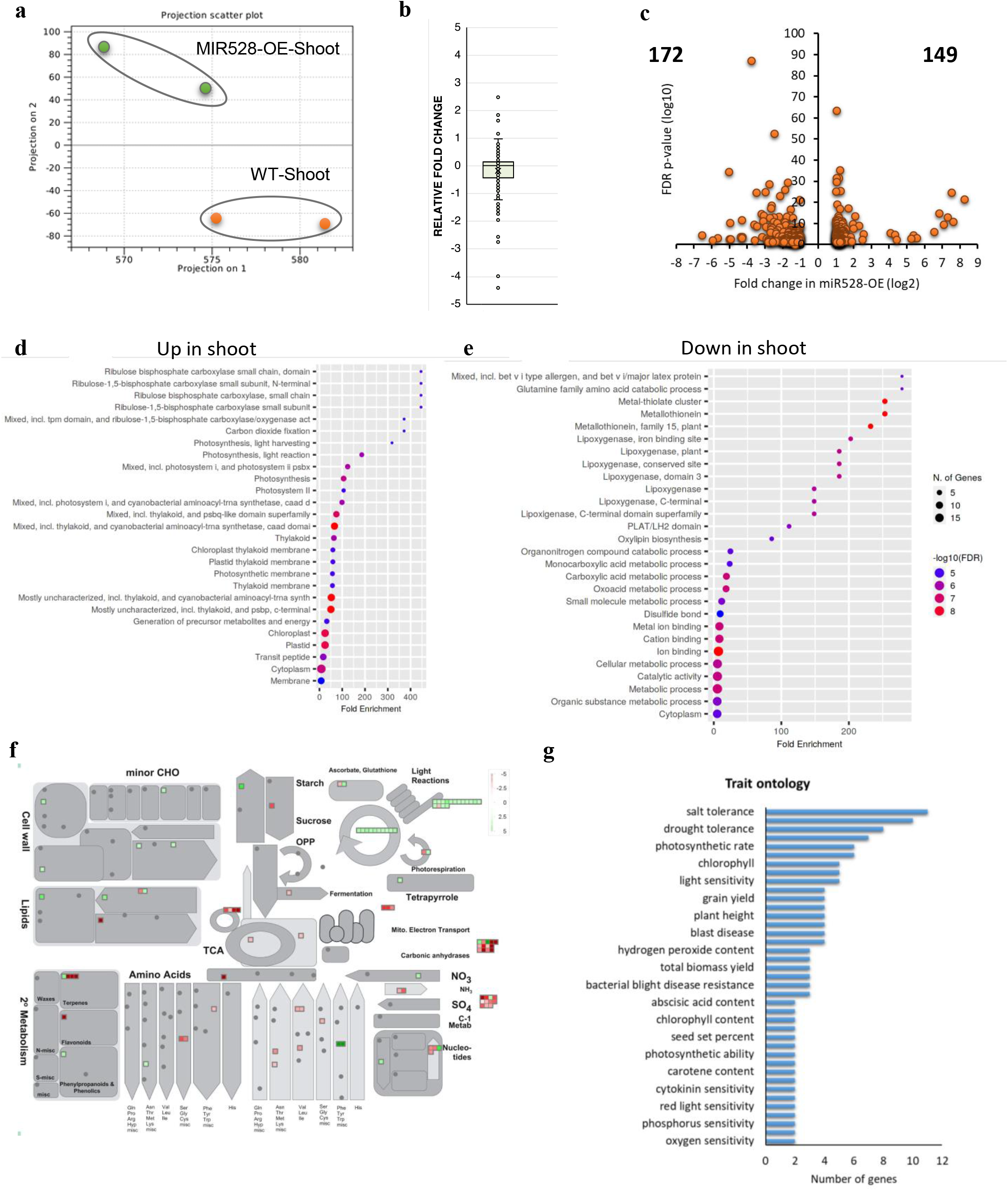
Transcriptome-wide regulation of miR528 in rice. **(a)** Principal component analysis of RNA-Seq data sets three week old WT and miR528-OE shoots. **(b)** The box plot showing the expression (relative fold expression) of miR528 target genes in shoots of MIR528-OE. **(c)** The differentially regulated genes in miR528-OE shoots are depicted in the scatter plot. The genes with up (FC≥2) and down regulated (FC≤-2) expression shown by plotting log10 of FDR pvalue versus log2 of relative fold change in miR528-OE w.r.t to WT. **(d-e)** Gene ontology enrichment analysis of differentially expressed genes. **(f)** The classification of DEGs using mapman showing the metabolic overview with significant upregulation of genes involved in photosynthesis. **(g**) The plots depicting the different traits associated with the DEGs.

### OsD3 is the species-specific target of miR528

In light of the observed tissue and drought-mediated expression dynamics of miR528/target modules and morphological and molecular traits exhibited by MIR528-OE plants, *OsD3* seems to be the most important target candidate in miR528-mediated regulations (Figure S8). Therefore, we went ahead to delineate the role of *the miR528:OsD3* node in rice SL signaling. *OsD3* encodes for the F-BOX/LRR-REPEAT MAX2 HOMOLOG which associates with OsD14, SL-receptor, and performs the degradation of SL-repressor protein, D53 allowing the downstream SL-mediated transcriptional responses via IPA1 and TRP proteins (Bürger & Chory, 2020a; Seto et al., 2019; Shabek et al., 2018). OsD3 is post-transcriptionally regulated by miR528 as clearly shown by the t-plots generated using the publically available degradome data, confirmation from previous studies (Figure 5a). It also exhibited a drought-mediated inverse expression pattern to miR528 in the root, flag leaf, and inflorescence but not at the tissue level under control conditions (Figure b-c). The qRT-PCR further validated the inverse expression dynamics of *miR528* and *OsD3* in roots and shoots of MIR528-OE seedlings (Figure 5d). The comparative transcriptomics approach was adopted to see the extent of shared transcriptional regulation of genes between RNA seq of *osd3* mutant (DEGs taken from Zheng et al. 2020) and MIR528-OE (present study) in rice. Several genes followed similar expression in *osd3* and MIR528-OE suggesting that *miR528*-mediated transcriptional modulations are partly regulated by a change in D3 levels. 118 upregulated (∼22.47%) and 322 downregulated (20.39%) genes shared similar expression in *osd3* and miR528-OE (Figure 5e-f) w.r.t respective WT plants. *miR528* is a monocot-specific miRNA while D3 is a highly conserved gene in the plant kingdom (Figure 5g). Considering the above fact, we enquire whether the miR528:D3 node is species/lineage-specific or not. It could be possible that the *miR528* target site might be conserved in plants but due to the absence of *miR528*, Os*D3* is not regulated by miRNA in plants. Therefore we extracted all the orthologs of D3 from the phytozome (https://phytozome-next.jgi.doe.gov/) and the cDNA of *OsD3* orthologs and miR528 mature sequence as input in PsRNAtarget (https://www.zhaolab.org/psRNATarget/) tool. Surprisingly only *OsD3* is predicted to be targeted with a significant alignment score (2) and other orthologs are targeted with a score beyond 5 (Figure 5h). Moreover, among the monocots also, low scores were obtained only in rice and not in others (Figure 5j). The above data suggest that *OsD3* is a species-specific target of miR528. Within the clade of Oryza, the target site of miR528 is conserved in *O. sativa, O. indica, O. nivara, O. rufipogon, O. barthii* and O. *meridionalis* while 1 and 3 mismatches were observed for *O. punctata* and *O. brachyantha* respectively (Figure 5i).

**Figure 5.**
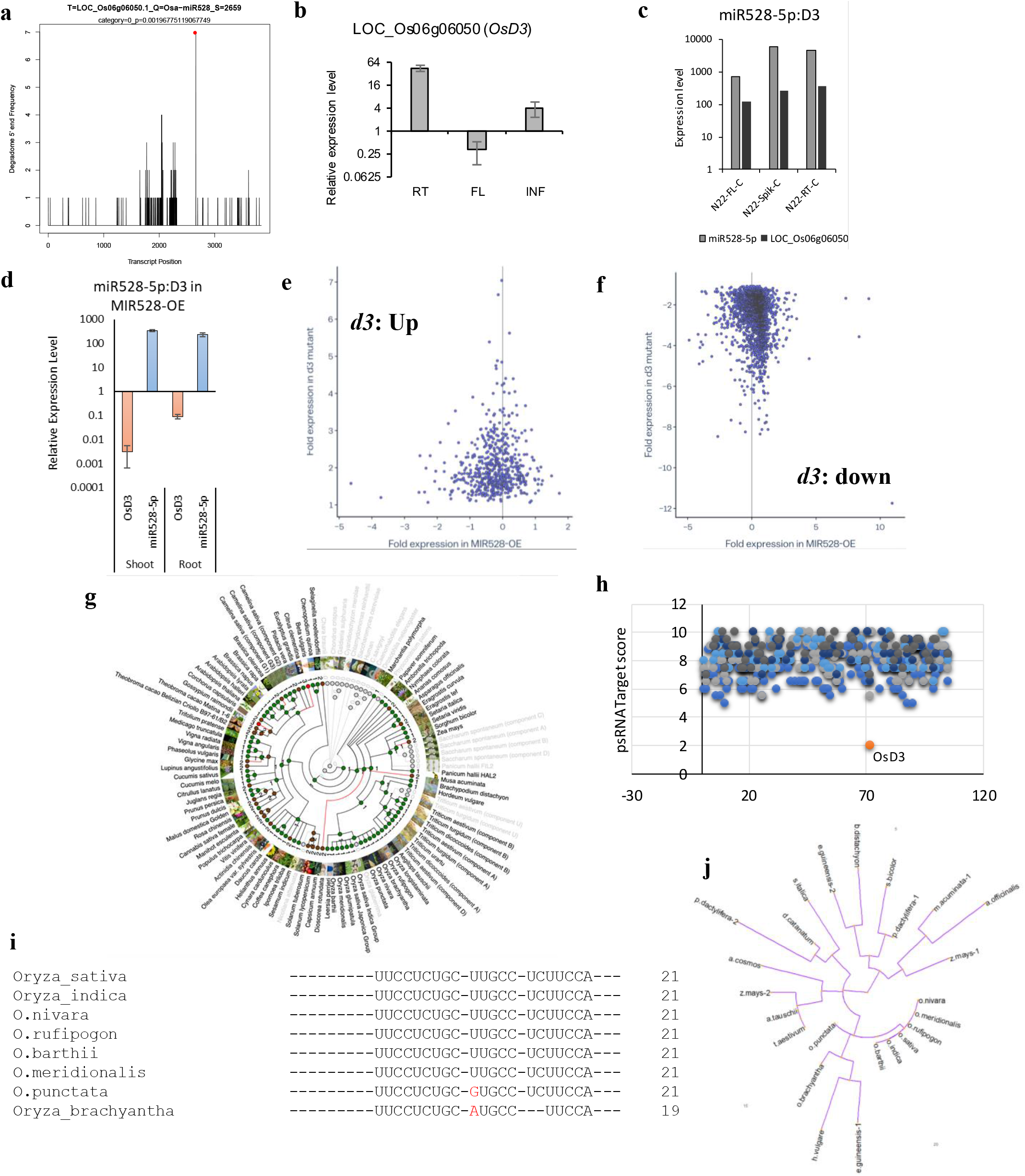
*OsD3* is *Oryza*-specific target of miR528. **(a)** T-plot of *OsD3* gene validating the cleavage by miR528. **(b)** The expression analysis of OsD3 in different tissues under drought stress using qRT-PCR. The values represent the mean of at least three biological and three technical replicate. Actin was used as the endogenous control. **(c)** Tissue mediated expression of miR528 and OsD3 in different tissues. **(d)** expression analysis of miR528 and OsD3 in shoots and roots of miR528-OE using qRT-PCR. 5s and actin was used as the endogenous controls for miR528 and OsD3 expression analysis respectively. Three technical and three biological replicates were used for calculation and error bars represents the standard error. **(e-f)** The scatter plot showing the comparative expression FC of DEGs {reported upregulated **(e)** and downregulated **(f**)} in *Osd3* mutant (Zheng et al. 2020) and miR528-OE. **(g)** The evolutionary conservation of OsD3 in plant kingdom. **(h)** The plot depicting the alignment score of miR528 on osD3 of different plant species. cDNA sequence of orthologs of D3 were used in target prediction of miR528. OsD3 score is highlighted in orange color. **(i)** Sequence comparison of miR528 target site in Oryza. **(j)** The evolution of miR528 target site in monocots based on the sequence variation.

### miR528 impacts the SL signaling via the post-transcriptional regulation of OsD3

No miRNA has been reported to regulate the SL biosynthesis and signalling in plants and regulated by SL itself in rice. As reported here also that *OsD3* is the high-confidence target of *miR528* but the interaction of this miRNA: target module with the SL signaling and its downstream effects are not clear. Therefore, to identify miRNAs targeting the genes involved in SL biosynthesis and signaling, the degradome data available at the pmiRen database is explored. As a result, miR1846a,f,h/D14, miR1852:D53 in addition to miR528/D3 regulatory modules have resulted (Table S5). But the expression of miR1846 and miR1852 was not detected in our small RNA datasets. So only the miR528:D3 module seems to be critical for in-depth analysis. In N22, all the SL biogenesis genes have very low expression while the signaling genes have a significant expression in flag leaf, inflorescence, and roots (Figure 6a). Moreover, higher expression was observed in roots and inflorescence as compared to the flag leaf (Figure 6a). *miR528* and *OsD3* showed tissue-specific differential expression under drought. As the complex of D14 and D3 is critical in determining the levels of SL-repressor protein D53 via proteasomal degradation and thereby regulates the downstream transcriptional network of SL response. Recently *Ideal Plant Architecture 1 (IPA1)*, a key protein involved in regulating the rice plant architecture including tillering and root development traits, was identified as the direct target of D53 (Song et al., 2017). Not only D53 interact with the IPA1 and suppresses its transcriptional activity but it is directly transcriptionally regulated by IPA1 forming the feedback regulatory loop (Song et al., 2017). Therefore, a decrease in *OsD3* levels should be reflected in D53 as well as SPL14 regulon. Interestingly in flag leaf where *miR528* was upregulated and *D3* was downregulated, *D53* was also downregulated while both *D3* and *D53* showed upregulation in roots inversely to that of miR528 levels (Figure 6b). The above observation suggests that the miR528:D3:D53 module was active under drought conditions. When the levels of SL biosynthesis and signaling genes were checked in miR528-OE plants, most of the genes (*CCD7, D14, FC1, TPR3*) including *D3* and *D53* showed downregulation (Figure 6c). The above-obtained expression was further validated through qRT-PCR in shoots of miR528-OE seedlings (Figure 6d). In miR528-OE as *OsD3* levels are decreased due to the post-transcriptional activity of *miR528*, we speculate that might lead to the transcriptional disturbance of IPA1-regulated genes. Therefore, to understand the impact of the miR528:D3 node on the IPA1-regulated transcriptome (Lu et al., 2013) at a global scale, the comparative transcriptome approach was adopted. The expression of IPA1 regulated genes was checked in miR528-OE and surprisingly the upregulated regulon of SLP14 (597 reported genes showing upregulation in SPL14 overexpression) exhibited a decline in expression pattern (308 genes with decreased expression in miR528-OE) while the IPA1-mediated downregulated regulon (665 genes showing downregulation in IPA1 overexpression) followed upregulation trend (229 followed elevated expression in miR528-OE) in miR528-OE shoots (Figure 6e-f). Moreover, the IPA1-target genes with enrichment of *GTAC* and *TGGGCC/T* motif in their promoter also followed an inverse trend in miR528-OE (Figure 6g-h). The known targets of IPA1 genes identified in the literature also exhibited decreased expression in miR528-OE as compared to WT shoots (Figure 6j). The above-identified miR528:D3/SPL14:D53 module was validated through qRT-PCR in miR528-OE plants (Figure 6i). The transcript level of *IPA1* was slightly upregulated in miR528-OE while both *D3* and *D53* exhibited downregulation, suggesting the involvement of protein level regulation of SPL14 via *D53* levels due to alteration in *D3* The above results clearly showed that the change in miR528 is playing important role in SL signaling via *D3* regulation.

**Figure 6.**
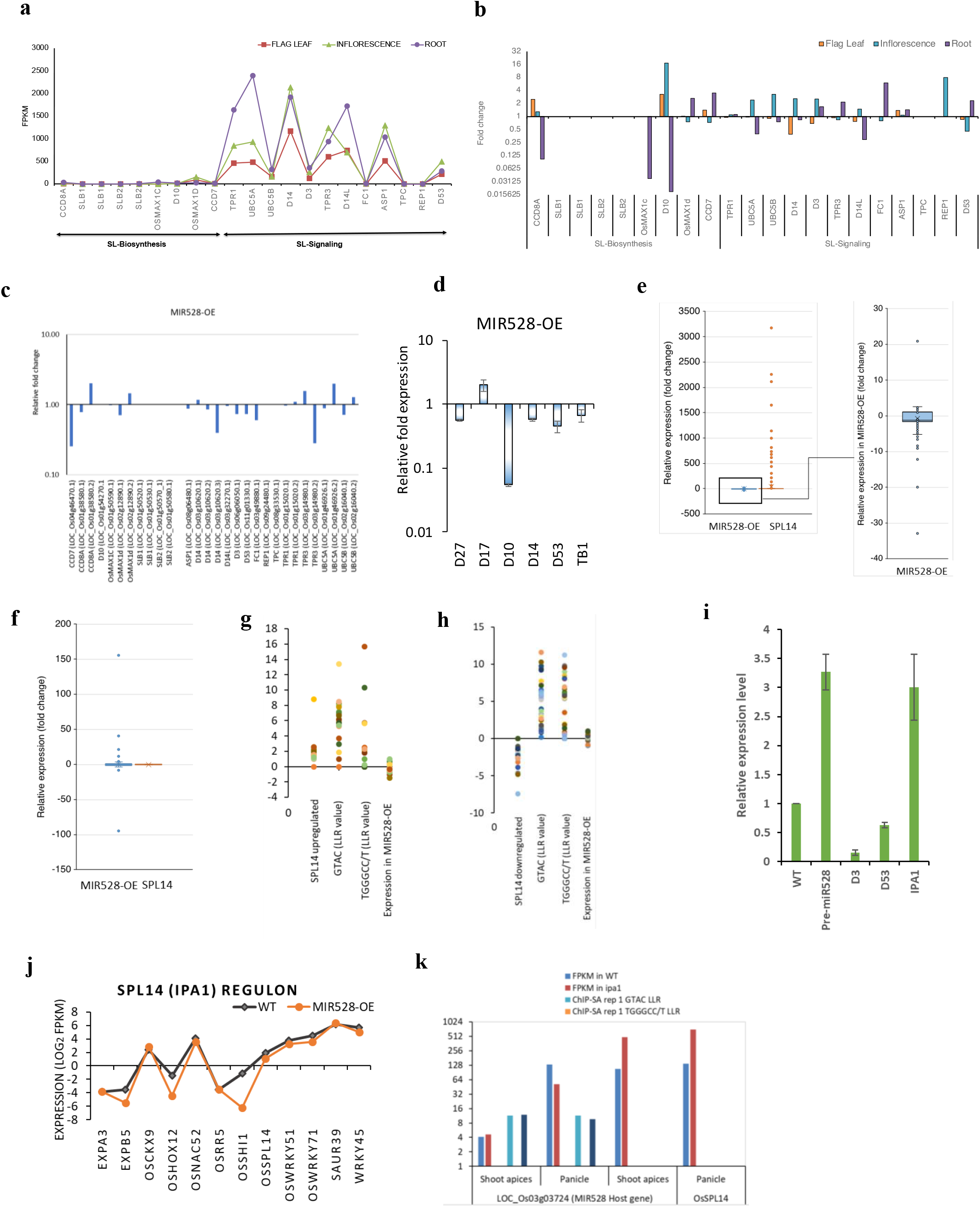
miR528 influence the SL signaling in rice. **(a-c)** The expression analysis of SL biosynthesis and signaling genes in different tissues of rice under control **(a),** drought stress **(b)** and miR528-OE shoots (**c**). **(d)** qRT-PCR expression analysis of some genes involved in SL biosynthesis and signaling. (**e-f)** comparative transcriptome of miR528-OE and DEGs of OsIPA1-OE demonstrating the opposite expression trend for IPA1 up (e) and downregulated **(f)** genes in miR528-OE. **(g-h)** the comparison of expression of IPA1 regulated (with IPA1 binding support) up **(g)** and downregulated **(h)** targets in miR528-OE transcriptome. **(i)** qRT-PCR validation of miR528:D3:IPA1:D53 module in miR528-OE shoot bases at seedling stage. (j) the comparative expression of IPA1 known targets in WT and miR528-OE plants. **(k)** evidence for the binding and opposite regulation of miR528 host gene in pull-down and transcriptome of IPA1.

### miR528:D3/SPL14:D53 module is also responsive to Nitrogen starvation

In rice, nitrogen starvation leads to a decrease in plant height and an increase in seminal root length in WT as well as in *d3* mutant (Luo et al., 2018). But in comparison to WT plants, *d3* seedlings had shorter plant height and decreased root length (Luo et al., 2018). Also, the increase in seminal root length was reported under low nitrogen conditions but the extent of low nitrogen-induced enhancement of SR length was diminished in *the d3* mutant (Sun et al., 2014). To ascertain the effect of miR528 overexpression on the above trait under nitrogen starvation conditions, both WT and miR528-OE seedlings were grown under control and nitrogen-deficient solution. As expected the classic nitrogen deficiency-induced increase in seminal root length was observed for both WT as well as miR528-OE seedlings (Figure 7a-b). Further, the enhancement in root length was less in miR528-OE in comparison to WT aligning with the phenotype observed for *the d3* mutant by Luo et al., 2018 (Figure 7b). Moreover, root length was reduced in miR528-OE seedlings even under control conditions (Figure 7b). But contrasting results on shoot length were observed in WT and miR528-OE, wherein the shoot length was decreased in miR528-OE w.r.t WT plants under low nitrogen conditions (Figure 7c). The above-observed root traits are SL-dependent, as the levels of SLs increase in response to nitrogen starvation (Sun et al., 2014). OsD3 is the key player in the SL-mediated root responses under nitrogen starvation responses. We went ahead to compare the GR responsiveness between WT and miR528 seedlings. In response to GR, the LR density was sharply decreased in WT plants and moderately reduced under low nitrogen conditions which mimic the GR24 response (Figure 7f). In miR528-OE roots, the effect of GR on LR density was negligible (Figure 7f). Interestingly, under low nitrogen, enhancement in the LR length was prominent in miR528-OE roots (Figure 7f). In addition, miR528 overexpression leads to increased AR and SR density under control conditions (Figure 7d). The LR density was reduced while the SR density was slightly increased under low nitrogen conditions. The above observations suggest that miR528-OE mimics the insensitivity response observed in d3 mutant (Sun et al., 2019) in response to GR24 treatment and nitrogen starvation in shaping root architecture in rice. Further to see whether miR528 itself is responsive to exogenous levels of GR24 or not? The expression of miR528 and its precursor was studied in the effect of exogenously applied synthetic rac-GR24 (Chiralix), and the inhibitor of strigolactone biosynthesis TIS108 (Figure 7f). The rice seedlings of WT and miR528-OE were grown in a medium supplemented with 2 μM rac-GR24 and 5 μM TIS108 for two weeks and shoot bases were used for expression analysis of miR528 and its precursor (Figure 7f). In response rac-GR24, both miR528 and its precursor followed downregulation as compared to the control (Figure 7f). In contrast, the expression of mature and precursor miR528 increases in shoot bases of seedlings treated with SL biosynthesis inhibitor, TIS108. The above results confirmed the negative transcriptional regulation of miR528 in response to SL levels.

**Figure 7.**
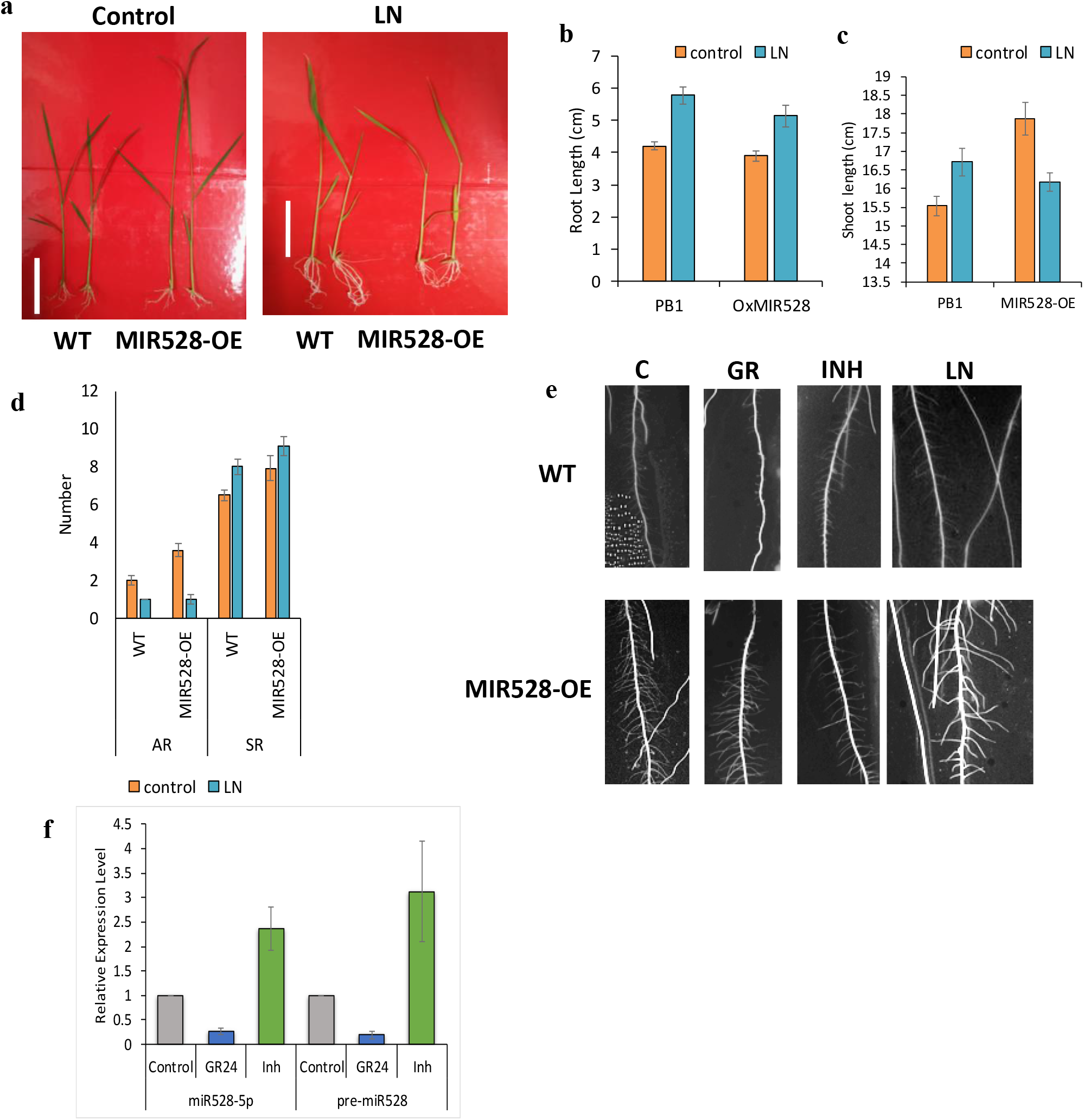
SL-insensitivity of miR528 plants. **(a)** Phenotype of WT and miR528-OE plants in response to low nitrogen (LN) conditions. (b-d) the measurement of root length **(b),** shoot length **(c),** AR and SR root density of WT and miR528-OE seedlings under control and LN conditions. **(e)** root phenotype of WT and miR528-OE plants under control **(C),** GR24 (GR), SL-inhibitor (INH) and low nitrogen (LN). **(f)** Expression analysis of miR528-5p and its precursor in response to GR24 and TIS108 in rice seedlings. Each value represents the mean of three biological and three technical replicates. Rice 5s and actin gene was used as the endogenous control for miRNA and gene expression respectively.

## Discussion

The results reported in the present study unfold the yet unexplored role of monocot-specific miR528 in regulating strigolactone signaling in rice. miR528 is only functional in monocots and shows high conservation among the different monocot families with single dominant mature sequences (41 monocot genomes analyzed here; Cuperus et al., 2011; Zhu et al., 2020). miR528 precursor sequences also exhibit a higher index of sequence similarity around the miR-miR* region including the consistent GC-rich and mismatch signatures consistent with observations by Zhu et al. 2020. In the present study, miR528 emerges as an interesting regulatory component as it exhibits tissue-biased drought response in the comparative analysis of miRNome of flag leaf, root, and spikelet in rice. In N22, miR528 is present in low levels in the flag leaf as compared to spikelet and root under control conditions, but under drought conditions the expression increases in the flag leaf and decreases in other tissues. Previously, the differential expression of miR528 is documented in response to heavy metals, H_2_O_2_, salt, nitrogen starvation, copper deficiency, and drought in rice. Moreover, it follows inverse expression in the flag leaf of drought-sensitive and tolerant cultivars of rice (Balyan et al., 2007). Being an exonic miRNA, it shares similar drought-responsive regulation with its host gene (LOC_Os03g03724) but the differs in tissue-level accumulation. While its tissue-biased expression correlates with its transcriptional regulator OsSPL9 in rice. The OsSPL9:miR528 module is involved in enhancing the antiviral defense and drought response in rice (Balyan et al., 2017.; Yao et al., 2019). The inspection of single base methylation levels in promoter and host gene region of miR528 in different tissues under control and drought suggested that methylation does not influence the transcriptional regulation of miR528. Further, the sequence comparison of miR528 precursor along with the regulatory region in 4726 rice accessions demonstrated that the precursor of miR528 is highly conserved with several SNPs in the promoter region coinciding with the cis-regulatory elements. The above observations suggest that transcriptional regulation is playing a critical role in maintaining the miR528 levels in response to developmental as well as environmental cues. miR528 act by regulating a diverse pool of target genes belonging to different gene families functioning in developmental to stress-responsive regulatory networks. The target pool of miRNA is influenced by several factors including the basal expression of miR-target gene, tissue types, and external conditions, etc. Further, it is possible that a miRNA inversely regulates one target, slightly alters the expression of another target, and not impact other targets at all. Therefore, the predicted target pool of miR528 was studied in detail in different tissue and drought conditions to identify the targets exhibiting anti-correlation in expression to miR528. In rice, 132 targets resulted from PsRNA prediction (score<5) and degradome data supports only 19 genes including *OsD3, OsAAO2*, and plastocyanin-like domain-containing proteins. Studies demonstrated the role of miR528 mediated regulatory roles of *OsAAO2, OsUCL23*, *OsD3*, and *OsRFI2* in antiviral response (Wu et al., 2017; Yao et al., 2019), pollen fertility (Zhang et al., 2020), plant height (Zhao et al. 2022) and flowering time (Yang et al., 2019) respectively. Through comparative expression analysis of *miR528* and its target transcripts by exploiting the miRNome and transcriptome of N22, only two targets {LOC_Os02g55600 expressed protein) and LOC_Os06g06050 (*OsD3*)} followed inverse expression pattern to *miR528* in all three tissues under drought. LOC_Os02g55600 is the predicted target only with a comparatively high score (5) as compared to *OsD3* which is a validated high-confidence target of *miR528*. *OsD3* encodes an F-BOX/LRR-REPEAT MAX2 HOMOLOG and is a critical component of the rice SL signaling pathway. In rice, SLs bind to DWARF14 (D14) which possesses both receptors as well as hydrolase activity and makes it compatible for OsD3 to interact through its dislodged C-terminal helix with D14 (Seto et al., 2019; Shabek et al., 2018). This D14-SCF(D3)-E3 complex targets the proteasome-mediated degradation of SL repressor, D53 and exerts SL-mediated transcriptional response via IPA1 and TPR proteins ( Zhou et al. 2013; Jiang et al. 2013; Song et al. 2017; Sun et al. 2021). As among the targets, OsD3 exhibited opposite regulation in all tissues in rice, it’s quite a possibility that it will affect the SL signaling and its downstream transcriptional events. Towards gaining insights on the role of *miR528:OsD3* in shaping SL signaling events, the detailed characterization of miR528 overexpression and its transcriptome were studied in light of the transcriptome of OsD3 mutant and OsSPL14 regulon. The following results confirm the involvement of miR528 in modulating the OsD3-mediated SL signaling pathway in rice:

1. miR528 overexpression exhibits many traits analogous to those reported for *Osd3* mutant including high tiller number, lateral root number, plant height (seedling stage), leaf width, chlorophyll content, and seminal root length.
2. *OsD3* appears to be an Oryza-specific target of miR528 since the evidence for targeting in other monocots is quite insignificant. The mRNA levels of *OsD3* showed a significant decrease in miR528 overexpression plants in addition to opposite expression to miR528 under drought in flag leaf, spikelet, and roots. Previous reports also validated *OsD3* as a genuine miR528 target (Balyan et al., 2017.; Zhang et al. 2020). At the transcriptome level also, 440 genes followed synergistic expression in the transcriptome of miR528 overexpression and *Osd3* mutant (Zheng et al., 2020).
3. Together with OsD14, OsD3 process the degradation of D53, SL repressor which associates with downstream targets such as TPRs (Jiang et al., 2013) and IPA1 (Song et al., 2017) and restricts their transcriptional activity. In overexpression, due to low *OsD3* levels, there would be increased restriction of D53 on its targets and subsequent repression of downstream transcription activity of its target transcription factor. The *miR528*-mediated downregulation was reflected in the repression in the transcriptional activation of D53 target IPA1. The comparative transcriptome of IPA1 overexpression (Lu et al., 2013) and miR528-OE highlighted an inverse correlation in the expression profile of 537 genes due to the decreased degradation of D53. IPA1 directly regulates the transcriptional activation of *OsD53* (Song et al., 2017), which also showed downregulated expression in miR528-OE plants.
4. Under low N conditions, miR528-OE demonstrated restricted seminal root elongation similar to the reported phenotype of *the Osd3* mutant (Luo et al., 2018) and d53 mutants (Sun et al. 2016). Further, in rice, *miR528* levels are induced in response to 2.5 mM nitrate levels and its overexpression exhibit enhanced biomass, increase root number, and nitrogen content under low nitrogen (Zhao et al., 2022).
5. miR528-OE mimics the insensitivity response observed in d3 mutant (Sun et al., 2019) in response to GR24 treatment and nitrogen starvation in shaping root architecture in rice.
6. Exogenous application of SL (GR24) negatively regulates *miR528* levels in rice.

In conclusion, the current study establishes a new regulatory node in the regulation of SL signaling in rice. This node is defined by miR528 which itself has been implicated in copper homeostasis, drought response, and plant development. What makes the association more interesting is that this regulatory node (miR528) appears to have evolved in a very specific manner in monocots only and behaves in a variety-specific manner under drought conditions. This uniqueness of miR528 imparts a very specific nature to the regulation of SL signaling in rice. The relevance of the evolution of such specific (rice-specific) regulation of SL signaling in plants needs to be further studied.

## Experimental Procedures

### Plant material and stress treatments

The *Oryza sativa* L. cv N22, was grown in fields, and drought was administered at the heading stage. Drought stress was measured by determining the soil moisture content (<15%) and visual scoring of leaf rolling phenotype. Flag leaf and spikelets (heading stage) and roots (milky stage) from control and stressed plants were collected and immediately frozen in liquid nitrogen and stored at −80L°C for further use.

For assaying the nitrogen starvation response of wild-type and miR528-OE plants, seeds were sterilized and maintained hydroponically in a rice growth medium (having 2.5 mM Nitrogen) under culture room conditions. Seven days old WT and miR528-OE seedlings of uniform growth were transferred to N deficient medium (0.02mM N) for two weeks. The seedlings were scored for morphometric parameters (seminal root length, shoot length, number of adventitious and seminal roots). For root phenotyping under N deficiency, rice seeds were surface sterilized and grown in vertically maintained square plates with solidified rice growth medium with sufficient and low N content. Plates were screened for root phenotype after 1 week of N deficiency.

To check the sensitivity of GR24 and complementation by TIS 108 on WT and miR528 OE seedlings. A synthetic rac-GR24 (Chiralix) was dissolved in anhydrous acetone according to the recommendations of Halouzka et al. (2020) to prepare 100 mM stock solution and used at 2 μM concentrations added to RGM medium. A strigolactone biosynthesis inhibitor, TIS108 (Chiralix), an effective tool for regulating strigolactone production in rice was dissolved in pure acetone before use to make a 100 mM stock solution further diluted to 5 μM final concentrations and added to RGM medium. Without RGM medium was used as a mock control for both GR24 and TIS108.The seedlings were grown hydroponically in growth chambers and cultured at 30°C under fluorescence white light with a 16 h light/8 h dark photoperiod for two weeks. 40 seedlings were taken for each treatment. After the treatment with 2 μM GR24 and 5 μM TIS shoot bases and root were collected and frozen at −80 °C until qRT-PCR analysis

### Small RNA sequencing

sRNA cDNA libraries from the mature roots of drought-tolerant N22 from control as well as drought-stressed plants were constructed using the Illumina small RNA v1.5 sample preparation kit as per manufacturer instructions and sequenced on the Illumina GAII sequencing platform. The raw sequencing reads were analyzed for their quality followed by the adapter (CAAGCAGAAGACGGCATACGA) removal using CLC genomics. The raw sequence reads were subjected to a computational analysis pipeline to remove the adapter sequences using the trim adapter tool of CLC genomics. Further adapter-cleaned reads were mapped to Rfam database 11.0 (Burge et al., 2012; http://rfam.xfam.org/) and the reads matching the structural RNA were removed. When the trimmed reads were mapped to their respective genomes (IRDB, Indica Rice Database, http://www.genomeindia.org/irdb/) a total of 16,855,251 (73%) sRNA tags mapped from N22 and PB1 libraries, respectively (Table 3.1). Further, reads that matched structural non-coding RNA (Rfam version 11.0, Burge et al., 2012; http://rfam.xfam.org/) corresponds to a value of 25%. To determine the profile of annotated miRNAs, the sRNA tags were mapped to the known rice miRNA genes in the ‘miRBase’ database (version 21; www.mirbase.org; (Kozomara & Griffiths-Jones, 2014) with the help of BLAST. The output was parsed with the help of in-house developed ‘perl’ scripts. The database had 592 precursors and 713 mature sequences (553 unique mature sequences) for Oryza sativa. About 2.86% of sRNA tags mapped to known miRNA genes. For comparative expression profiling based on sRNA tag counts the number of tags mapping to each miRNA gene was normalized to transcript per million (TPM). For comparative miRNome analysis, the sRNA profiling data of flag leaf and spikelets generated in our previous study (Balyan et al., 2017) were used. Principal component analysis and hierarchical clustering of small RNA datasets were done using CLC genomics workbench version 9.

### RNA-Seq analysis

The expression of miR528 targets and other genes discussed in the present study in flag leaf, spikelet, and roots of N22 under control and drought stress conditions were extracted from our published study by Guar et al., 2021. The list of DEGs reported in *Osd3* mutant and *OsIPA* was obtained from studies by Zheng et al. 2020 and Lu et al 2013 respectively. The expression data from the above studies are used for comparative transcriptomics.

For comparative transcriptome analysis of WT and miR528-OE, high-quality RNA of shoots of two-week-old seedlings of WT and miR528-OE were outsourced for paired-end RNA-Seq in two biological replicates on illumine platform. The quality control of raw RNA-Seq data was performed using the FastQC followed by trimming using TrimGalore. Good quality adapter trimmed reads were analyzed following the RNA-Seq tool of CLC genomics as per default parameters. High-quality adapter trimmed reads were analyzed by RNA-Seq plug-in of CLC genomics workbench version 9 using the rice genome and cDNA sequences from Rice Genome Annotation Project (RGAP version 7) as the reference on default parameters [mismatch cost: 2, insertion cost: 3, deletion cost: 3, length fraction: 0.9, global alignment: no, auto-detect pair distance: yes, strand specificity: both, the maximum number of hits for a read: 10, expression value: RPKM (Reads Per Kilobase of transcript per Million mapped reads)]. The expression data were analyzed by the ‘Empirical analysis of Differentially Expressed Genes (DEGs)’ algorithm of the CLC genomics workbench to perform the two-group comparison following exact test (Robinson et al., 2010) in the EdgeR Bioconductor package using default settings. Principal component analysis (PCA) of all the RNA-Seq data sets was performed using CLC genomics workbench version 9. The genes with dispersion fold change of ≥2 or ≤−2 with FDR correction P-value of ≤0.05 (WT vs. miR528-OE) were considered as differentially regulated by miR528. Gene ontology analysis of DEGs was performed using the ShinyGO tool (v 0.741 http://bioinformatics.sdstate.edu/go/) following a P-value cutoff (FDR) of ≤ 0.05. The DEGs were assigned to different metabolic pathways using MapMan Metabolic overview tool v-3.6.0RC1 (Thimm et al., 2004). Only those bins were considered whose p-value was ≤ 0.05. Trait ontology terms associated with the DEGs were extracted from the Oryzabase database (https://shigen.nig.ac.jp/rice/oryzabase/).

### Sequence Variation and Haplotype Evaluation of miR528 loci in 4726 Rice Accessions

The DNA sequence variations (SNPs and Indels) in the miR528 precursor, host gene, and its 1kb upstream regulatory regions were detected in 4726 accessions using the ‘variations by region tool’ of RiceVarMap v2.0 (http://ricevarmap.ncpgr.cn; (Zhao et al., 2021)). The variation IDs detected in the promoter region were then used as input for haplotype analysis using the ‘Haplotype Network analysis’ tool of RiceVarMap2.

### Determination of differential Methylated Cytosines (mCs) in miR528 loci

Methylation levels in different tissues under control and drought conditions were determined by analyzing the whole genome bisulfite datasets using the bisulfite sequencing plugin of the CLC Genomics Workbench (version: 9, Qiagen) following default parameters. Briefly, to call mCs, we deployed a bidirectional protocol to map to the reference genome. The mapped reads were used as an input for calling the methylation level based on a Fisher exact test (p-value ≤ 0.05) after removing non-specific matches, duplicate matches, and broken pairs read. The error rate was determined by aligning reads to rice chloroplast and found to be ∼1.44% in both the control and drought stress samples. After incorporating the error rate, binomial distribution was calculated and the mCs were filtered based on a p-value ≤ 0.005 with at least 5 reads in both the replicates. The mCs passing all these criteria were used for further analyses. Transcript information from RGAP (Rice Genome Annotation Project; V7.0; http://rice.uga.edu; Kawahara et al. 2013) was used to annotate the methylated sites through R Bioconductor packages of GenomicFeatures v 1.46.1 (Lawrence et al., 2013) and ChIPSeeker (v1.30; Yu et al. 2015). All the analyses were performed using custom R scripts built in-house. All the relevant information regarding methylated cytosines in the promoter and precursor region was extracted for flag leaf, roots, and inflorescence under drought conditions.

### Generation and characterization of miR528 over-expression rice plants

To generate the transgenic plants over-expressing miR528-5p, the binary vector pB4NU with maize ubiquitin promoter was selected. To generate the Ubi::MIR528 construct, the 88 bp (Chr3: 1667328-1667415 [+]) pre-miR528 was PCR amplified from genomic DNA isolated from N22 seedlings using Phusion taq DNA polymerase (Thermo) using the primers; 5’-AGCGGTACCAGTGGAAGGGGCATGCAG-3’ and 5’-AGAGCTCGGAATGGAAGAGGCAAGCA-3’. The 88 bp PCR product was gel eluted and digested with the KpnI and SacI. Similarly, the pB4NU is also digested with the same set of restriction enzymes. The KpnI and SacI digested 88bp MIR528 fragment and pB4NU was ligated using the T4 DNA ligase (Fermentas) as per the given directions. Then the 3 µl of ligation mix was transformed into the E.Coli XL1B cell using chemical transformation and plated. The construct cloned in E.coli was confirmed by colony PCR, digestion, and sequencing and then moved to Agrobacterium for plant transformation. The transgenic overexpression MIR528 under drought-sensitive PB1 background was generated as per the protocol of Toki et al. (2006) plants. The transgenic lines (T3 generation) were analyzed for their morphometric differences from the wild-type plants. The following traits were scored for both WT and miR528-OE; days to heading, number of tillers, flag leaf length and width, plant height, internode length, number of primary branches in panicle, number of grains per panicle, number of effective grains per panicle, seed length and width, chlorophyll fluorescence. The data were represented in the form of graphs. For most of the lines, more than 10 plants were analyzed for each parameter. For phenotyping, rice seeds from the WT plants and the miR528 OE transgenic plants were imbibed in darkness for 2 d at 30 °C and then were grown for ∼ 20 d in hydroponically at 28 °C, 70% humidity (16 h light/8 h dark), and then the seedlings were transplanted to a field at NIPGR. During this period, the average low-temperature range is ∼ 22.9 –25.5 °C, and the average high-temperature range is ∼29.7– 40 °C. Plants were maintained under routine management practices. The grain length, width and thickness, the number of primary and secondary branches, the number of effective grains, and the internode length were determined when the seeds were harvested. For each line, data from 10 or more individual plants were obtained and subjected to statistical analyses.

### Multiple Sequence Alignment and Phylogenetic Analysis of miR528 Family

The mature and precursor sequences of miR528 from monocots were extracted from miRbase (https://mirbase.org/), PmiREN2.0 (Plant miRNA Encyclopedia; https://www.pmiren.com; (Guo et al., 2022) and sRNAanno (http://plantsrnas.org/). Multiple sequence alignment followed by phylogenetic tree construction of precursor miR528 was performed using the alignment tool of CLC genomics (version: 9, Qiagen). The evolutionary tree was constructed using the maximum likelihood method and the Tamura-Nel model (Tamura et al., 2011).

### miRNA and mRNA Expression Analysis

The total RNA was extracted using TRI-reagent (Sigma-Aldrich) followed by DNAse I treatment (Thermo Scientific) as per the manufacturer’s instructions. The quality and quantity of RNA were analyzed on Nanovue (GE Healthcare Life Sciences) followed by verification of RNA (1 μg) on formaldehyde agarose gel (1.2 %). The high-quality total RNA (A260/280 = 1.8 – 2.0 and A260/230 = ≥ 2.0) was used to enrich the small RNA using equal volumes of 4M LiCl followed by polyadenylation using a Poly(A) Tailing Kit (Ambion). An amount of 2 μg of polyadenylated small RNA and total RNA was reverse transcribed using miR_oligodT_RTQ (100 bp) (Ro et al. 2006) and oligodT (25 bp), respectively, followed by cDNA synthesis using SuperScript II Reverse Transcriptase (Invitrogen) as per instructions. All the primers for the real-time PCR analysis were designed using the ‘Primer Express 2.0’ software (expected product size = 100–120 bp) (Life Technologies) followed by the individual pair confirmation using the BLAST program in the RGAP database (rice.plantbiology.msu.edu; Kawahara et al., 2013) MicroRNAs (miRNAs) a) against genomic and cDNA sequences.

To analyze the expression of miRNAs, qRT-PCR was performed using a reaction mixture (7 μl) containing cDNA (1 μl), 2X Taqman Fast Universal PCR master mix (3.5 μl, Applied Biosystems) with 10 mM miR408 specific forward primer (0.6 μl), 10 mM universal fluorogenic probe (0.16 μl) specific to miR_oligodT_RTQ and 10 mM RTQ universal reverse primer (0.6 μl) in Rotor-Gene Q (Qiagene) according to the manufacturer’s protocol. For target gene expression and pre-miR408, Fast SYBR Green master mix (3.5 μl) was used with gene-specific primers (0.3 μl each), and 1 μl of the diluted cDNA template to make the final reaction of 7 μl was run on Rotor-Gene Q (Qiagene) as per manufacturer’s protocol. Rice 5S and Actin were used as endogenous control for miRNA and mRNA expression, respectively. The ΔΔCt method was followed to calculate the relative fold change (2^−ΔΔCt^) in expression. The sequence of primers is listed in Table S7.

### Chlorophyll Content and Fluorescence Measurement

Chlorophyll was extracted from the wild type (WT) and transgenic leaves with dimethyl sulfoxide (DMSO). An amount of 50 mg of finely chopped leaves was incubated with 1 ml of DMSO at 65 °C for about 1 h. A volume of 50 μl of the extract was diluted to 200 μl and the absorbance was measured at 645 nm and 663 nm on infinite® M200 PRO (Tecan). The concentration of total chlorophyll and chlorophyll a/b was calculated by the equation proposed by Arnon (1956). The chlorophyll fluorescence parameters were measured using a Pulse-amplitude modulation fluorometer (Junior-PAM, H. Waltz, Germany). Plants were dark-adapted for 30 min before measuring the maximum photosynthetic efficiency (Fv/Fm). The electron transport rate (ETR) and effective photosynthetic efficiency (YII) were recorded in light-adapted plants. More than 10 plants were used for each reading.

## Author Contributions

SR conceived the research and supervised the experiments; SB and DS executed all the experiments and analyzed the data; SK and RJ analyzed the methylation status of the miR528 region; SK performed the phylogenetic analysis of miR528 target site in D3 orthologs; VP and TAC performed qRT-PCR analysis of selected targets; RC complemented the expression analysis; SR and SB wrote the manuscript.

## Short legend for supporting information

**Table S1.** Summary of small RNA sequencing data

**Table S2.** The expression of miRNA in rice roots under control and drought stressed conditions. The values represent the TPM levels.

**Table S3.** The involvement of drought regulated miRNA targeted genes in different pathways of rice. The analysis was performed using the analysis tool of reactome (https://reactome.org/).

**Table S4.** The list of miR528 targets

**Table S5.** The expression levels (mean values) genes obtained in transcriptome of wild type and miR528-OE plants. The expression value represents the mean of two biological replicate.

**Table S6** The miRNA targeting the genes involved in SL and KR biosynthesis. The rice degradome data was obtained from pmiREN database.

**Table S7.** List of primer sequence.

**Figure S1.** Trait and gene ontology analysis of DEGs obtained in roots under drought.

**Figure S2.** The evolutionary conservation and divergence of miR528s in plant kingdom.

**Figure S3.** Comparative analysis of methylation levels in miR528 host gene and its upstream region.

**Figure S4.** Ontology enrichment analysis of genes targeted by miR528-5p.

**Figure S5.** Sequence variants and haplotype analysis of miR528 and its promoter region in 4726 rice accessions.

**Figure S6.** Construction of miR528 overexpression in rice

## Funding statement

This work was funded by the Department of Biotechnology (DBT), Government of India (Grant no. BT/PR3311/AGR/2/817/2011).

## Conflict of interest

The authors declare that the research was conducted in the absence of any commercial or financial relationships that could be construed as potential conflict of interest.

## Notes

### Competing Interest Statement

The authors have declared no competing interest.

