## supplemental data for "Monocot-specific *miR528* act as the post-transcriptional regulator of strigolactone signaling via *Dwarf 3* in rice"

[1] Equal authorship

* **Corresponding author:**

Prof. Saurabh Raghuvanshi

Department of Plant Molecular Biology,

University of Delhi South Campus, New Delhi 110021, India;

Orcid ID: 0000-0002-8349-4290

**All author Affiliations:**

Department of Plant Molecular Biology, University of Delhi, South Campus

**Email:** Sonia Balyan; Deepika Sharma, Shivani Kansal, Vaishali Panwar, Ringyao Jajo, Tonu Angaila Chithung and Saurabh Raghuvanshi

**Table S1**. Summary of small RNA sequencing data

| sRNA Data | Root Control | Root drought |
|---|---|---|
| Raw Reads | 2,44,91,133 | 2,84,55,433 |
| No. of reads after trimming | 2,26,66,487 | 1,64,65,197 |
| Genome matched | 1,28,65,281 | 90,01,827 |
| Reads mapped to rfam | 20,29,143 | 37,65,744 |
| Matched miRBase | 3,14,659 | 85,172 |
| Unique mature miRNA matches | 272 | 259 |

**Table S3**. The involvement of drought regulated miRNA targeted genes in different pathways of rice. The analysis was performed using the analysis tool of reactome (https://reactome.org/).

| miRNA | Target | Description | Pathway |
|---|---|---|---|
| osa-miR156k | LOC_Os03g28330.3 | sucrose synthase, putative, expressed | Galactose degradation II |
| osa-miR156k | LOC_Os06g05060.1 | ELF3 protein, putative, expressed | Circadian rhythm |
| osa-miR156k | LOC_Os08g02700.1 | fructose-bisphospate aldolase isozyme, putative, expressed | Calvin cycle |
| osa-miR156k/osa-miR156l-5p | LOC_Os03g03720.1 | glyceraldehyde-3-phosphate dehydrogenase, putative, expressed | Plastid glycolysis |
| osa-miR156k/osa-miR156l-5p | LOC_Os06g44970.1 | auxin efflux carrier component, putative, expressed | Root elongation |
| osa-miR156k/osa-miR156l-5p | LOC_Os07g32170.1 | OsSPL13 - SBP-box gene family member, expressed | Regulation of seed size |
| osa-miR156k/osa-miR156l-5p | LOC_Os08g39890.1 | OsSPL14 - SBP-box gene family member, expressed | Response to submergence |
| osa-miR156k/osa-miR156l-5p/osa-miR156l-5p | LOC_Os04g55920.1 | zinc-finger protein, putative, expressed | Jasmonic acid signaling |
| * osa-miR164e | LOC_Os02g08440.2 | WRKY71, expressed | Jasmonic acid signaling |
| * osa-miR164e | LOC_Os12g08810.1 | VTC2, putative, expressed | Ascorbate biosynthesis |
| osa-miR167h-3p | LOC_Os03g14540.1 | UDP-glucuronate 4-epimerase, putative, expressed | GDP-D-rhamnose biosynthesis |
| osa-miR167h-3p | LOC_Os04g37619.1 | zeaxanthin epoxidase, chloroplast precursor, putative, expressed | Carotenoid biosynthesis |
| osa-miR171h | LOC_Os09g38030.1 | UTP--glucose-1-phosphate uridylyltransferase, putative, expressed | Galactose degradation II |
| osa-miR1850.1 | LOC_Os07g10770.1 | CESA8 - cellulose synthase, expressed | Cellulose biosynthesis |
| osa-miR3979-3p | LOC_Os03g22810.1 | copper/zinc superoxide dismutase, putative, expressed | Removal of superoxide radicals |
| osa-miR3979-3p | LOC_Os09g31478.2 | auxin efflux carrier component, putative, expressed | Intracellular auxin transport |
| osa-miR437 | LOC_Os01g14610.2 | PSF2 - Putative GINS complex subunit, expressed | DNA replication Initiation |
| osa-miR437 | LOC_Os02g05700.2 | OsFBO8 - F-box and other domain containing protein, expressed | Circadian rhythm |
| osa-miR437 | LOC_Os04g46960.4 | glutathione peroxidase domain containing protein, expressed | Glutathione redox reactions I |
| osa-miR528-5p | LOC_Os02g54890.1 | UDP-glucuronate 4-epimerase, putative, expressed | GDP-D-rhamnose biosynthesis |
| osa-miR528-5p | LOC_Os06g06050.1 | OsFBL27 - F-box domain and LRR containing protein, expressed | Strigolactone signaling |
| osa-miR530-5p | LOC_Os12g13320.1 | argininosuccinate synthase, chloroplast precursor, putative, expressed | Citrulline-nitric oxide cycle |
| osa-miR530-5p | LOC_Os12g13320.1 | argininosuccinate synthase, chloroplast precursor, putative, expressed | Arginine biosynthesis |

**Table S6.** The miRNA targeting the genes involved in SL biosynthesis and signalling. The rice degradome data was obtained from pmiREN database.

| miRNA | Target ID | Target Description | PsRNATarget Score | Degradome Category | Degradome datasets |
|---|---|---|---|---|---|
| Osa-miR1846a | LOC_Os03g10620 | D14 | 4 | 2 | SRR039716,SRR039718,SRR1609325,SRR1849887,SRR1849888,SRR1849889,SRR1849890,SRR1849892,SRR1849893,SRR1849894,SRR1849902,SRR1849903,SRR1849904,SRR1849905,SRR1849906,SRR1849907,SRR1849908,SRR1849909,SRR1849910 |
| Osa-miR1846f | | | 4 | 2 | SRR1609325 |
| Osa-miR1846h | | | 4 | 2 | SRR3140959,SRR3140960 |
| Osa-miR528 | LOC_Os06g06050 | D3 | 2 | 0 | SRR039716,SRR039717,SRR039718,SRR039719,SRR039720,SRR1609324,SRR1609325,SRR3140958,SRR3140959,SRR3140960 |
| Osa-miR1852 | LOC_Os11g01330 | D53 | 4 | 2 | SRR039718,SRR3140958 |

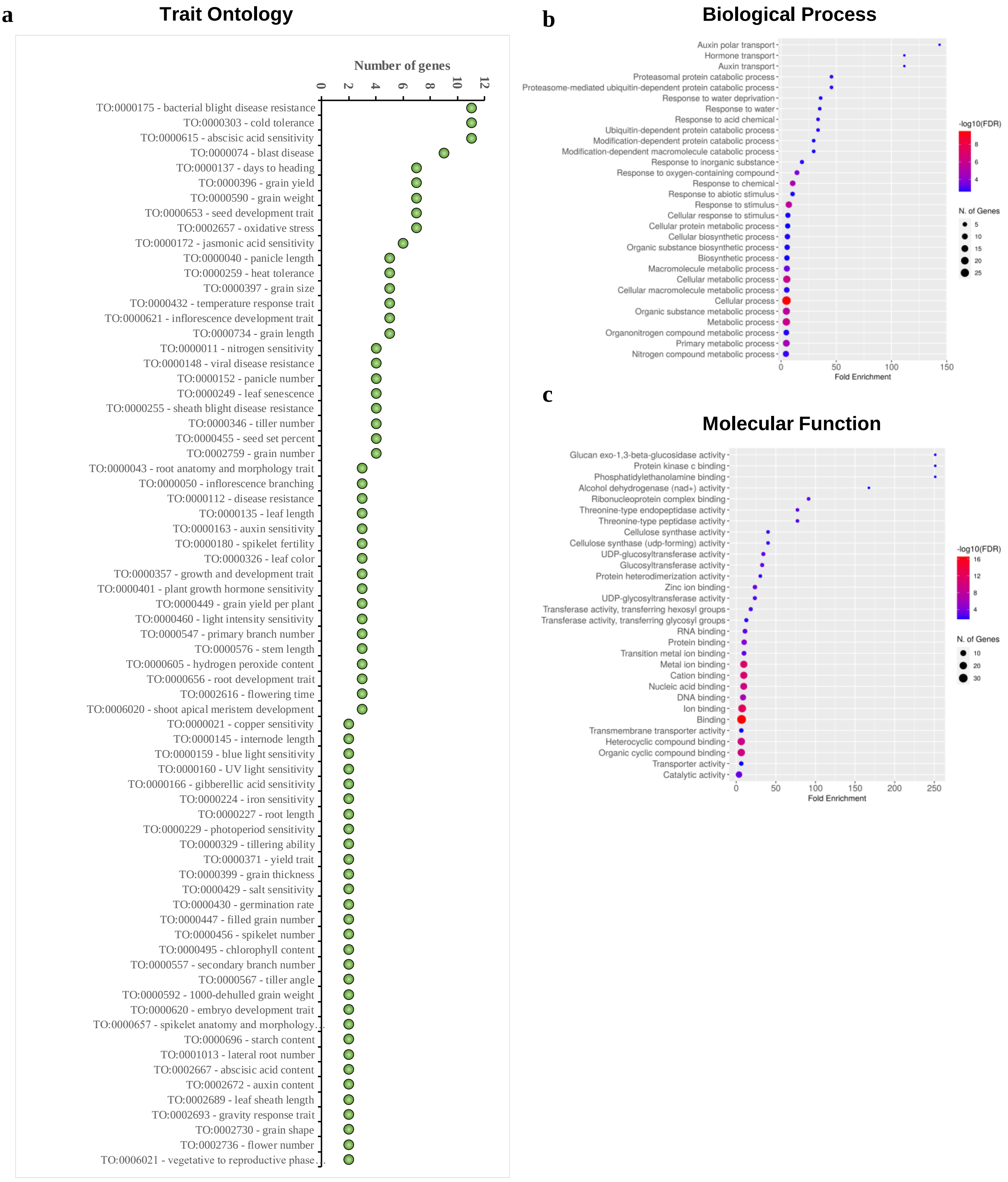


**Figure S1**. Trait and gene ontology analysis of DEGs obtained in roots under drought. The trait (a) and GO-enrichment analysis (b-c) of genes regulated by differentially expressed drought regulated miRNAs in roots of N22 using Oryzabase and ShinyGO tool respectively.

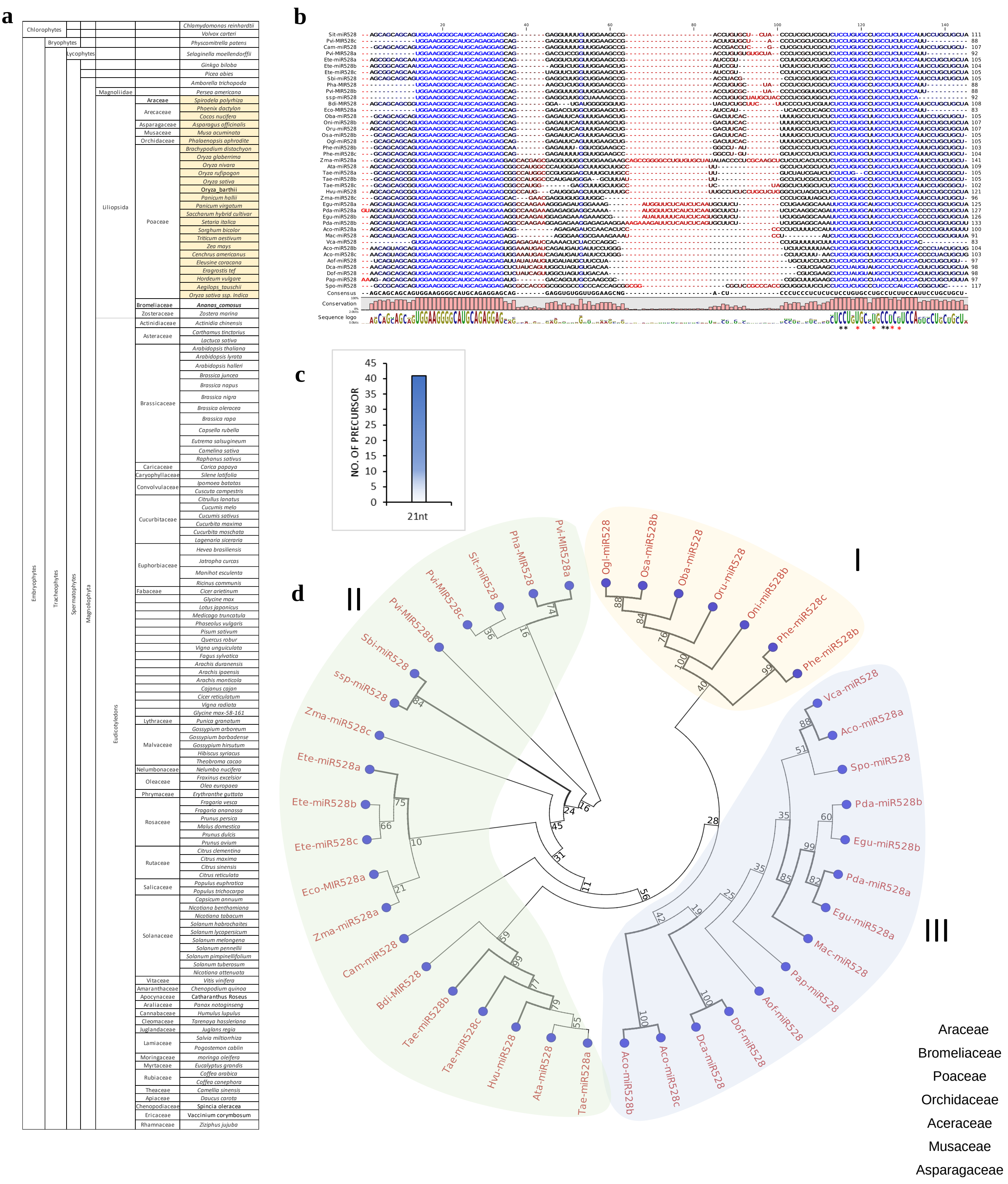


**Figure S2. The evolutionary conservation and divergence of miR528s in plant kingdom**. **(a)** The chart demonstrating the presence of annotated miR528 in plant species obtained from different public small RNA repositories. **(b)** Sequence comparison of miR528 precursors across liliopsida. The multiple sequence alignment tool of CLC genomics workbench was used. **(c)** plot showing the size distribution of mature miR528s in across Liliopsida. **(d)** The phylogenetic analysis of precursor miR528 in plants. The evolutionary tree was constructed using the maximum likelihood method and Tamura-Nel model (Tamura et al. 2021).

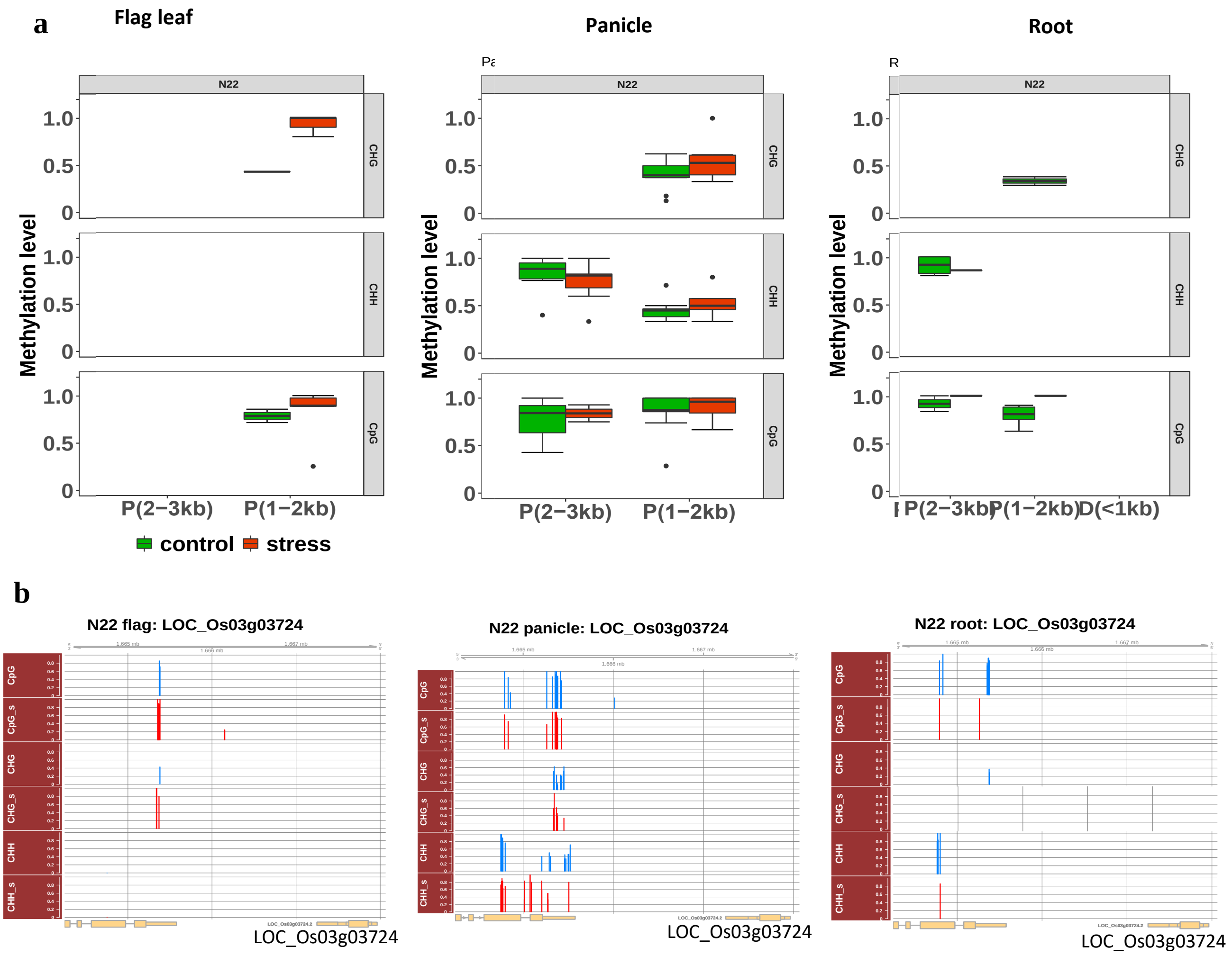


**Figure S3. Comparative analysis of methylation levels in miR528 host gene and its upstream region**. **(a)** Box-plots showing the methylation levels of different context i.e. CpG, CHH and CHG in flag leaf, spikelet and roots of N22 under control and stress conditions. **(b)** Localization of methylation peaks *w.r.t* to LOC_Os03g03724. Only the methylation levels following the p-value criterion of ≤ 0.05 were shown in the Figure.

a

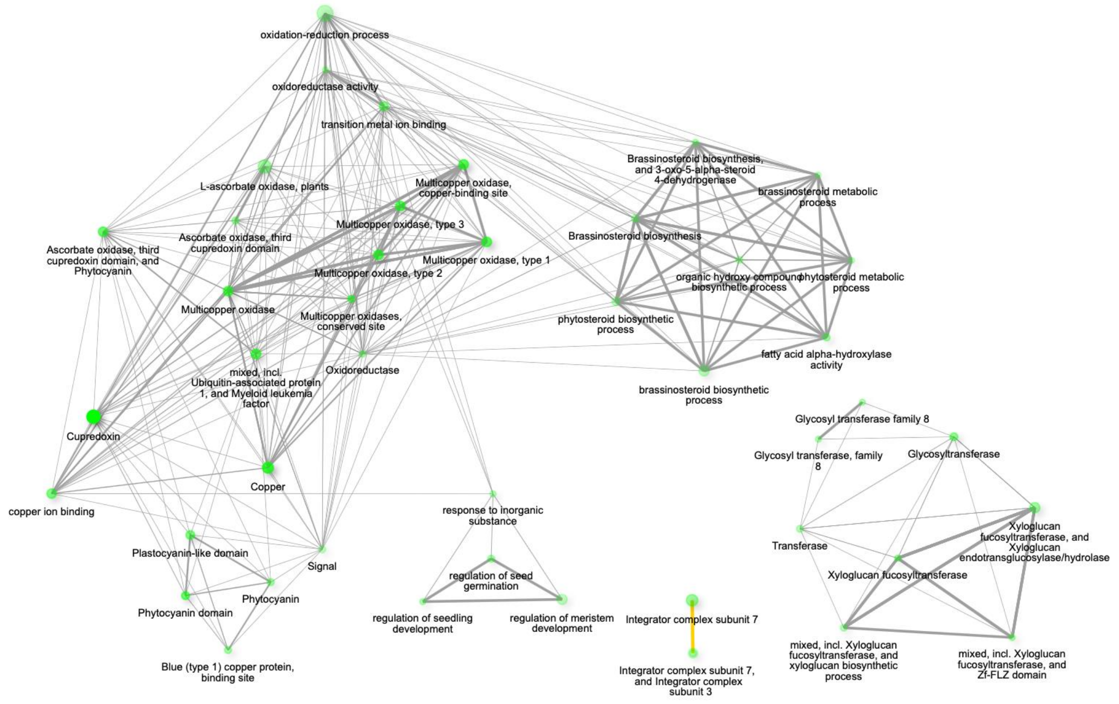


b

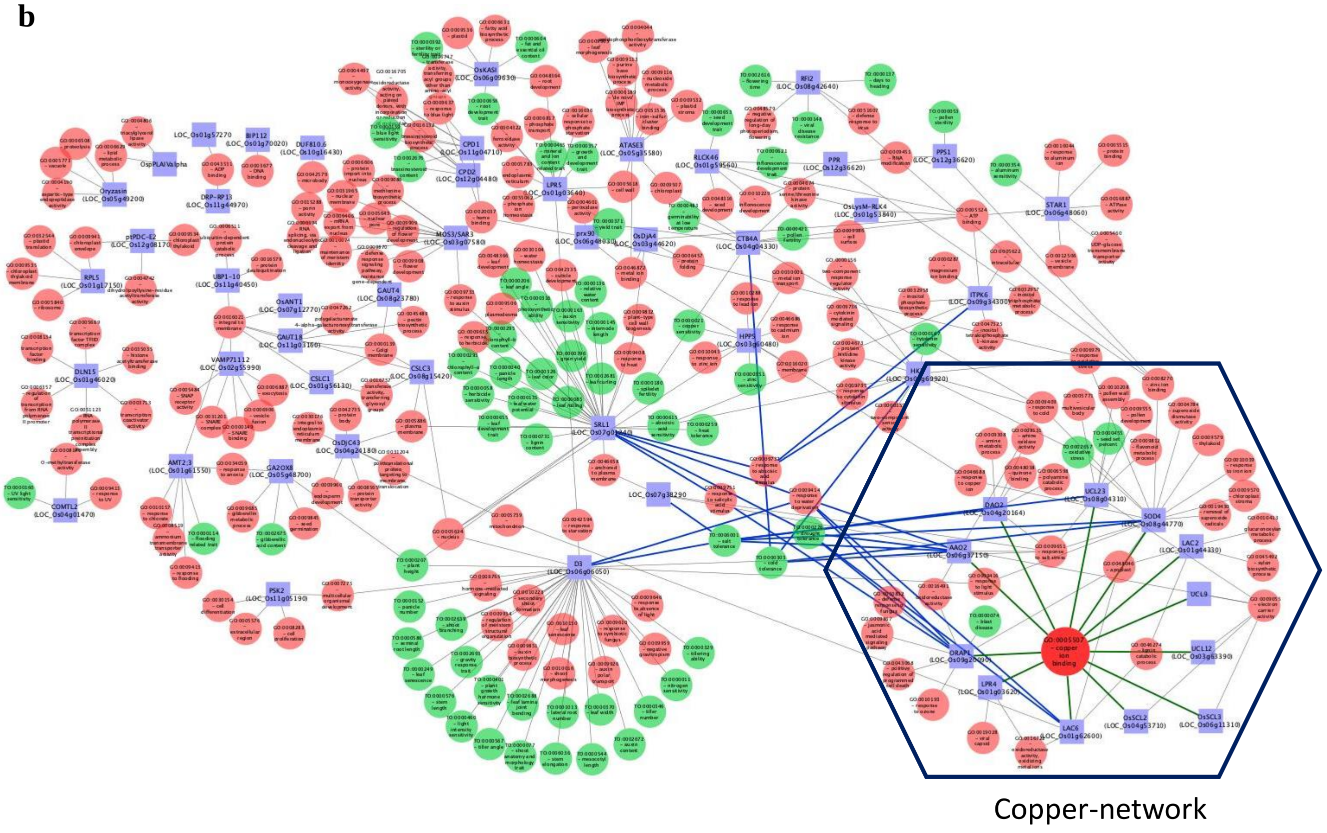


**Figure S4. Ontology enrichment analysis of genes targeted by miR528-5p**. GO enrichment analysis **(a)** and network showing the GO and trait ontology terms associated with genes targeted by miR528-5p **(b)**. Target, GO and TO are represented by blue square, red and green circle respectively. Targets associated with copper binding were highlighted in the section.

a

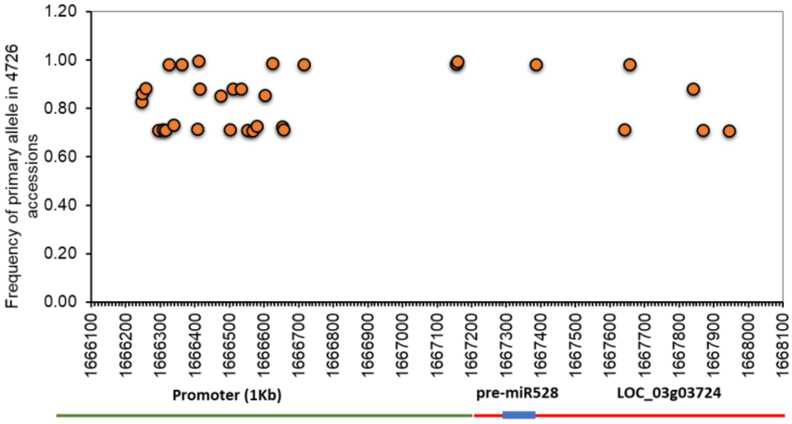


b

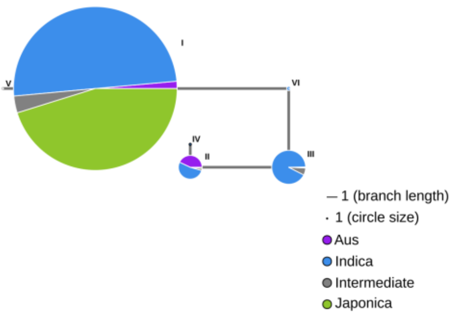


c

| Haplotype Group | Haplotype | ALL Frequencies | Indica | Japonica | Intermediate |
|---|---|---|---|---|---|
| I | ACACTGGTCTAGTAAACGTGGC | 3293 | 1695 | 1488 | 110 |
| II | ATGACGATCTGACGGACGCGCG | 477 | 456 | 7 | 14 |
| III | GCGACGATCAGGTGGGCGCGGG | 682 | 631 | 5 | 46 |
| IV | ATGACGATCTGACGGACACGCG | 43 | 43 | 0 | 0 |
| V | ACACTGGTCTAGTAAATGTGGC | 20 | 19 | 0 | 1 |
| VI | NCGACAGCCTGGTGGACGCAGG | 82 | 81 | 0 | 1 |

d

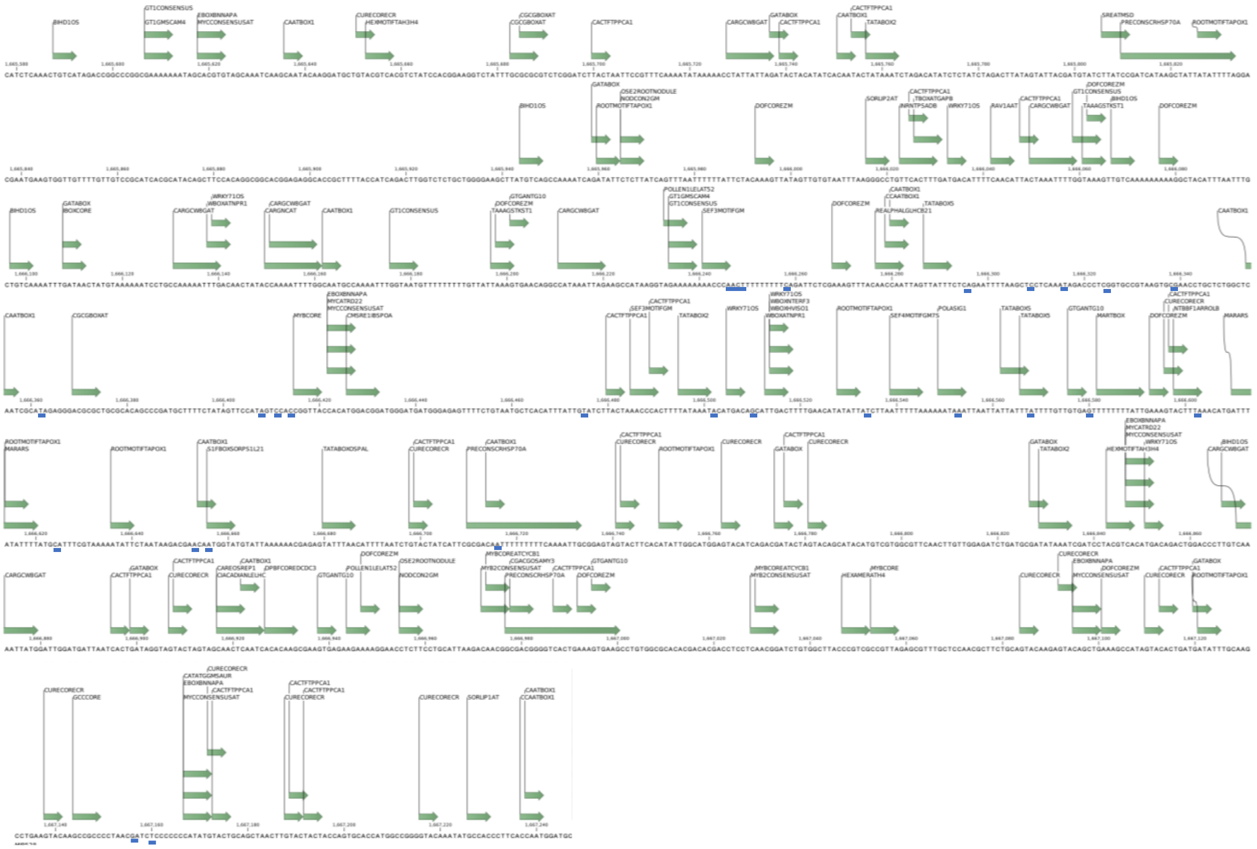

**Figure S5. Sequence variants and haplotype analysis of miR528 and its promoter region in 4726 rice accessions.** **(a)** The plots show the position and frequency of the primary allele of variants identified in the region of host gene of miR528 and its promoter region (1 kb) using the RiceVarMap2 database. The coordinates marked with green, blue and red lines represent the promoter, precursor miR528 and host gene (LOC_Os03gg03724) respectively. **(b)** The plot shows the result of the haplotype network analysis generated by RiceVarmap2. Only haplotypes found in ≥ 10 rice accessions were used to construct the haplotype network. **(c)** The details of six haplotypes were detected on the basis of variants in the promoter region. **(d)** Mapping of variants and cis-regulatory elements on the promoter region. The motifs were identified using New place online tool.

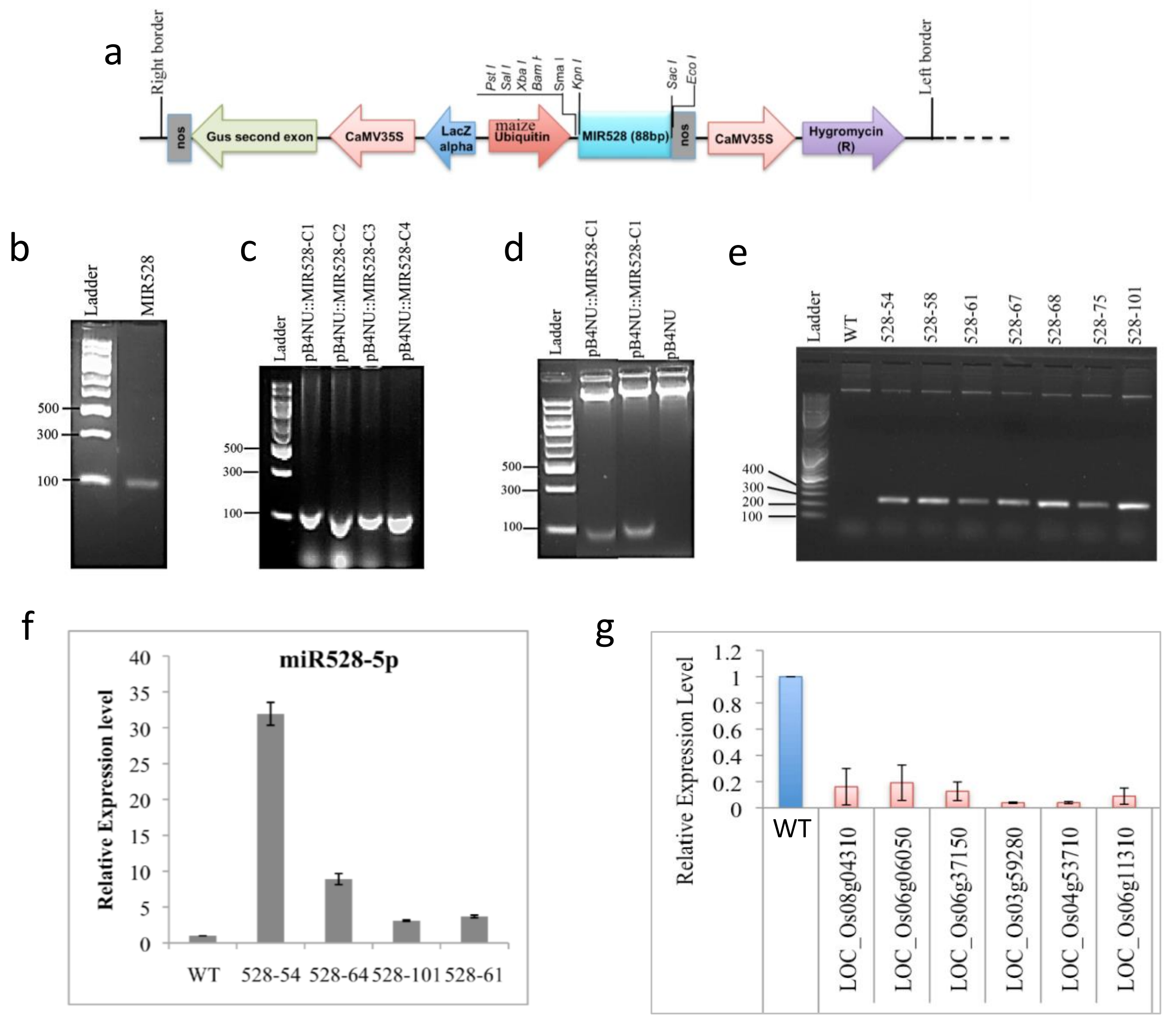


**Figure S6 Construction of miR528 overexpression in rice (a)** Cloning strategy of MIR528 in binary vector pB4NU. **(b)** PCR amplification of precursor miR528. Confirmation of recombinant clones harboring pB4NU::MIR528 by colony PCR **(c)** and digestion with KpnI and SacI **(d).** PCR confirmation of rice transgenic lines harboring pB4NU::MIR528 using maize ubiquitin specific primers **(e). (f)** Confirmation of transgenic lines by expression profiling of miR528-5p in different lines obtained. 5S is used as the endogenous control and WT seedlings were used as control. Three technical and three biological replicates were used. **(g)** The expression analysis of some miR538 target genes using qRT-PCR in line 64. Rice actine gene is used as the endogenous control and WT seedlings were used as control. Three technical and three biological replicates were used for calculation and error bars represents the standard error.

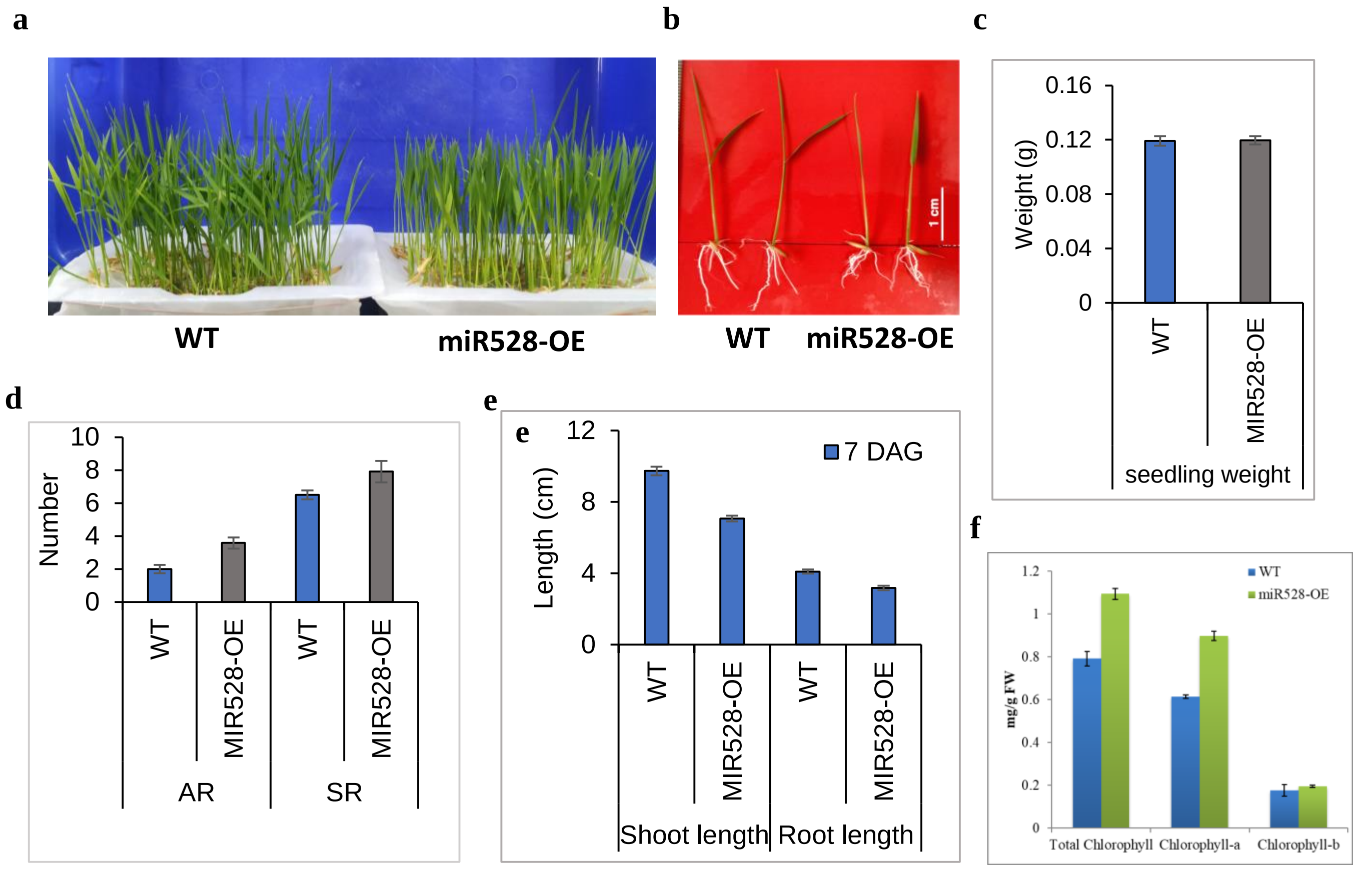


**Figure S7. Morphometric analysis of WT and miR528-OE at seedling stage. (a-b)** one-week old seedlings of WT and miR528-OE. **(c-e)** Comparative analysis of seedling weight, root density and shoot and root length of WT and miR528_OE seedlings. (f) Chlorophyll content in wild type and transgenic lines in seedlings.

### D3 (LOC_Os06g06050.1)

- **Morphological traits**
- Leaf senescence
- Stem length
- Tiller number
- Lateral root number
- Mesocotyl length
- Leaf width
- Plant height
- Tiller angle
- Gravity response
- Shoot branching
- Panicle number
- Seminal root length
- Tillering ability

- **Environmental response**
- Salt tolerance
- Nitrogen sensitivity
- Light sensitivity
- Phosphorus sensitivity
- Response to strigolactone

- **Molecular response**
- Transcriptional regulation of D53
- Altered transcriptional regulon of SPL14
- Altered transcriptional regulon of SPL14
- Transcriptional regulation of downstream genes of SL signaling

### common

- **Morphological traits**
- Shoot length
- Tiller number
- Lateral root number
- Panicle number
- Seminal root length
- Seminal root number

- **Environmental response**
- Nitrogen sensitivity
- Phosphorus sensitivity
- response to strigolactone

- **Molecular response**
- Trancriptional regulation of D53
- Altered transcriptional regulon of SPL14
- Transcriptional regulation of downstream genes of SL signaling

### MIR528-OE

- **Morphological traits**
- Stem length
- Tiller number
- Lateral root number
- Leaf width
- Plant height
- Panicle number
- Seminal root length
- Tillering ability
- Seminal root number
- Days to heading
- Flag leaf length
- Panicle branching
- Grains per panicle
- Seed length

- **Environmental response**
- Drought response
- Nitrogen sensitivity
- Copper sensitivity
- Phosphorus sensitivity
- response to strigolactone
- Calcium sensitivity

- **Molecular response**
- Trancriptional regulation of D53
- Altered transcriptional regulon of SPL14
- Photosynthetic rate
- Transcriptional regulation of downstream genes of SL signaling

**Figure S8**. The comparison of traits observed (miR528) and reported (Osd3) for miR528:D3 module

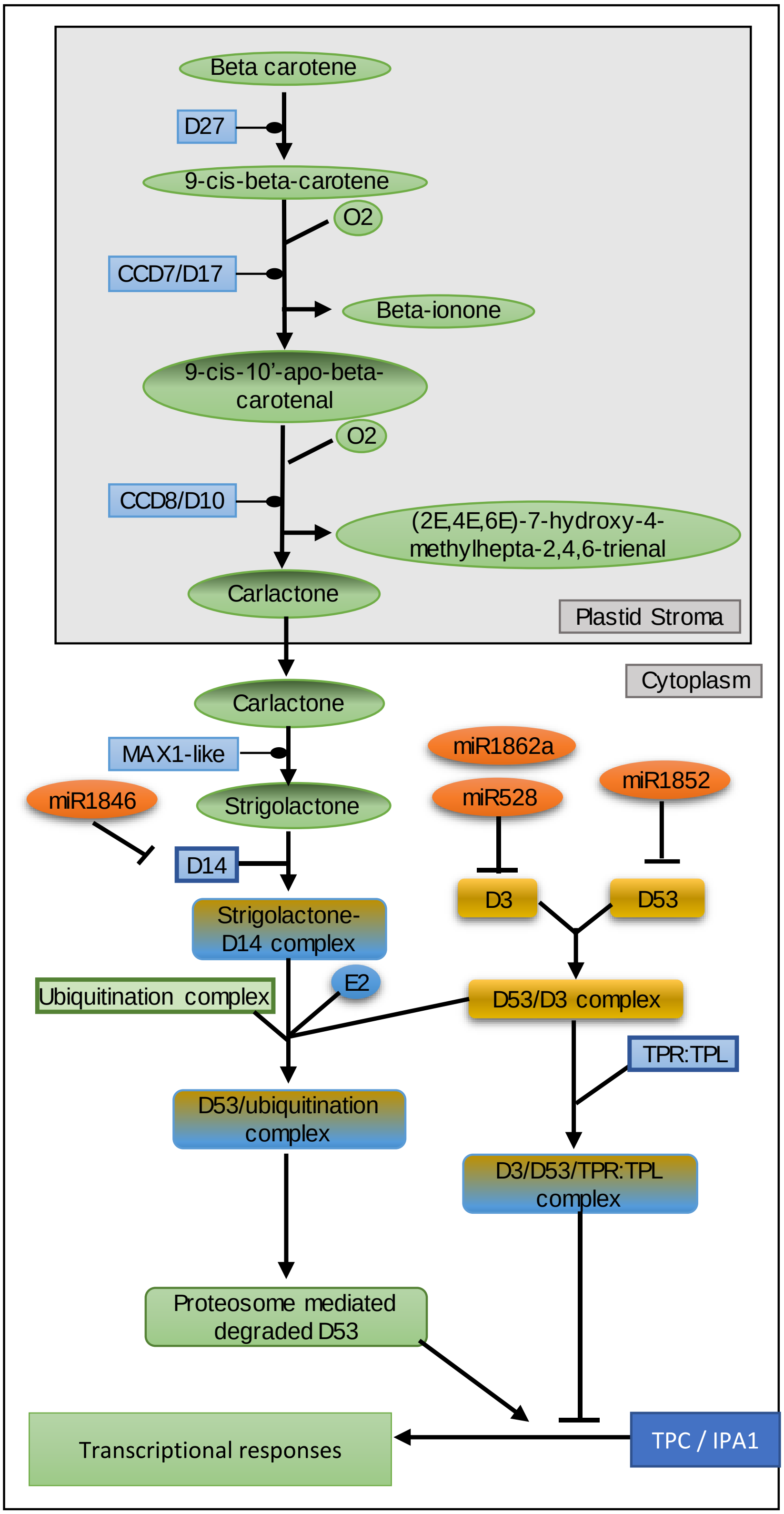


**Figure S8.** Diagram depicting the strigolactone biosynthesis and signaling in rice
